# Exploring the phylogenetic diversity and antimicrobial activity of non-*aureus staphylococci* and *mammaliicocci* isolated from teat apices of organic dairy cows

**DOI:** 10.1101/2024.02.01.578391

**Authors:** F. Peña-Mosca, T. N. Gaire, C. Dean, P. Ferm, D. Manriquez, P. Pinedo, N. Noyes, L. Caixeta

## Abstract

Prior studies have suggested that non-*aureus staphylococci* and *mammaliicoci* (**NASM**) possess inhibitory activity against mastitis pathogens. However, their impact on udder health outcomes and the mechanisms underlying this potential protective effect remain poorly understood. Our first objective was to identify NASM species on teat apices of organic dairy cows, assess their within-species phylogeny, and explore associations with presence of intramammary infections (**IMI**) and genomic features, including antimicrobial peptides (**AMPs**), virulence, and resistance genes. The second objective was to evaluate the *in vitro* antimicrobial activity of NASM isolates against mastitis pathogens and examine its associations with taxonomy, phylogeny, AMP genes, and IMI. Milk and teat apex swabs were collected weekly from 114 cows on two organic farms. Milk was cultured to identify *Staphylococcus aureus* (**SAU**) or *Streptococcus* spp. and *Streptococcus*-like organisms (**SSLO**) IMI. A case-control was designed to include cows with and without SAU or SSLO IMI. For each selected cow, the teat apex gauze swab collected during the week preceding IMI diagnosis (or corresponding time for controls) was aerobically cultured, and the taxonomy of isolates was determined using mass spectrometry. Isolates classified as NASM were subjected to whole genome sequencing using Illumina MiSeq. The inhibitory activity of NASM isolates was tested against SAU and *Streptococcus uberis*. Phylogenetic trees were constructed using Snippy and IQ-TREE. Genomes were assembled and annotated to identify species, AMP genes, virulence, and antimicrobial resistance markers. The *in vitro* antimicrobial activity of NASM varied across species and between cows with and without an IMI. *Staphylococcus succinus* was the species most frequently associated with highly inhibitory isolates, which were more prevalent in cows without IMI (19.4% vs 5.8%). Organic dairy cow teat apices harbored multiple NASM species and strains. All isolates had at least 1 AMP associated gene in their genome. *In vitro* antimicrobial activity was generally unrelated to clade membership, except for isolates classified as *Staphylococcus succinus*. *Staphylococcus aureus* had high virulence gene prevalence, while NASM species showed lower, species-specific prevalence. This study advances understanding of NASM antimicrobial activity and virulence potential.

## INTRODUCTION

Mastitis is the leading cause of morbidity (USDA, 2007) and milk production loss in dairy farms (Heikkila et al., 2018). Although preventive measures are important for the control of mastitis, antimicrobials are still a major component of on-farm contagious mastitis control (Ruegg, 2017). However, the availability of antimicrobials for dairy farmers and veterinarians is at-risk due to concerns about antimicrobial resistance to medically important antibiotics and policy restrictions on antimicrobial use in livestock (McCubbin et al., 2022; Rajala-Schultz et al., 2021; Ruegg, 2022). This limitation is especially remarkable on United States organic-certified dairy farms, where regulations for production of organic milk include a strict prohibition against the use of antimicrobials and other synthetic substances (NMC, 2019).

Prior data from our group showed that *Staphylococcus aureus* (**SAU**) and *Streptococcus* spp. had a high prevalence and persistence in the mammary gland of dairy cows in organic farms in the first 35 days of lactation (Peña-Mosca et al., 2023). Moreover, this study showed that SAU and *Streptococcus* spp. in early lactation were associated with a higher risk of high somatic cell count (**SCC**) during the first 180 days of lactation in organic dairy cows (Peña-Mosca et al., 2024). Non-*aureus staphylococci* and *mammaliicocci* (**NASM**) are also highly prevalent in milk samples (De Buck et al., 2021; Peña-Mosca et al., 2023), but unlike SAU and *Streptococcus*, in some studies, NASM IMI have been associated with increased milk production (Compton et al., 2007a; Piepers et al., 2010); while others suggested no association between NASM IMI and milk production (Nyman et al., 2018; Tomazi et al., 2015; Valckenier et al., 2020). The protective effects of NASM, have been supported by results from experimental studies in which the infusion into the mammary gland of NASM has been associated with decreased risk for IMI with SAU and *Streptococcus* spp., suggesting a potential protective effect of NASM against these two pathogens (Reyher et al., 2012). *In vitro* studies using isolates obtained from the mammary gland of dairy cows (i.e., from milk or teat apex) have shown that NASM are capable of inhibiting the growth of SAU (Carson et al., 2017; De Vliegher et al., 2004a; Toledo-Silva et al., 2022). Using the cross-streak method for inhibitory testing, these studies reported that between 9.1% and 14.3% of the isolates were able to inhibit the growth of SAU (Carson et al., 2017; Toledo-Silva et al., 2022). Recently, agar dilution methods have also been utilized to investigate the inhibitory capabilities of NASM species on mastitis pathogens (Toledo-Silva et al., 2022). One potential explanation for the *in vitro* antimicrobial activities is the ability of NASM to produce antimicrobial peptides (**AMPs**) (Braem et al., 2014; Carson et al., 2017; Nascimento et al., 2005), commonly referred to as bacteriocins or *Staphylococcins* (de Freire Bastos et al., 2020; Newstead et al., 2020). While AMP-encoding genes on bovine NASM isolates have been explored (Braem et al., 2014; Carson et al., 2017; Nascimento et al., 2005), further research is needed to identify specific AMPs that could be targeted for development of products for mastitis control. Lastly, although there has been significant expansion in our understanding of NASM epidemiology in recent years, the number of studies that have comprehensively investigated the entire genome of NASM are still limited (De Buck et al., 2021; Fergestad et al., 2021; Naushad et al., 2019). This limits our ability to investigate associations between NASM genotypes, phenotypes, and cow-level udder health outcomes. Therefore, more research is needed to describe the genomic diversity of NASM isolates from dairy cows, including important phenotypic drivers such as genes related to AMPs, virulence factors and antimicrobial resistance genes. Understanding the presence and diversity of these genes could provide further insights into the potential for NASM isolates to act as protective commensals or, conversely, as emerging pathogens in the udder environment.

Our first objective was to identify the NASM species present on teat apices of organic dairy cows and examine their within-species phylogeny, as well as explore associations with intramammary infections (**IMI**) and genomic features, including genes related to AMPs, virulence, and antimicrobial resistance. The second objective was to assess the *in vitro* antimicrobial activity of NASM isolates cultured from teat apices of cows with and without IMI, and explore its associations with NASM taxonomy, within-species phylogeny, the presence of AMP genes, and IMI.

## MATERIALS AND METHODS

This manuscript was prepared following the guidelines outlined in the Strengthening the Reporting of Observational Studies in Epidemiology - Veterinary (STROBE-Vet) Statement (Sargeant et al. in 2016). All study activities were approved by the University of Minnesota Institutional Animal Care and Use Committee (IACUC) (Protocol Number: 1807: 36109A), and Colorado State University IACUC (Protocol number: 1442). Laboratory activities were approved by the University of Minnesota Institutional Biosafety Committee (IBC) protocol: 2205-40001H.

### Inclusion criteria

Only cows from USDA organic certified dairy farms were enrolled in this study. Farms from different regions of the United States and with different herd sizes were selected. Enrollment was based on willingness to participate, and proximity to the Universities involved in the study. This work was part of a larger research initiative to investigate potential associations between the udder microbiome and udder health (Dean et al., 2021). Following the study design to assess that objective, cows from 5 organic dairy farms were enrolled 8 weeks before calving and followed up during the first 5 weeks of lactation (Peña-Mosca et al., 2023). In this manuscript, data from only two of these farms were included, as teat apex samples for bacteriological culture were only collected in these two farms due to limited research labor availability. Specifically, this manuscript used samples collected from a subset of 114 cows (Farm A [n =21] and Farm B [n =93]) out of the 596 cows enrolled in the main research project, where both milk and teat apex samples were collected. Samples from Farm A were collected between August 2019 and January 2020 and included all first-lactation cows that calved during that period. Samples from Farm B were collected between February and July 2021 and included both first and second lactation cows that calved throughout that interval.

### Milk sample collection

After calving, quarter milk samples were collected weekly during the first 5 postpartum weeks prior to the morning milking by members of the research team, following guidelines from the National Mastitis Council (NMC, 2017). Upon removal of the pre-dipping disinfectant, 3 to 4 streams of milk were discarded from each quarter, and the teat apex of each quarter was scrubbed with a gauze soaked in 70% ethanol. Milk samples were collected in a clockwise direction, beginning with the right rear quarter, and ending with the right front quarter. Approximately 10 mL of milk from each quarter were collected into separate 60 mL plastic vials (Capitol Vials, Thermo Fisher Scientific). Samples were placed on ice immediately after collection and stored in a freezer at -20 °C until further processing.

### Milk pooling

Quarter milk samples were combined into composite samples using the following protocol (Peña-Mosca et al., 2023). First, milk samples were thawed at 4 °C overnight. Next, quarter milk samples were homogenized by inverting them upside down twice. Then, 2 mL was extracted from each vial and dispensed into a single plastic vial inside a laminar hood. Lab-pooled composite samples were then submitted to the Laboratory for Udder Health at the University of Minnesota (St. Paul, MN, USA) for milk culture.

### Milk culture

Using a cotton swab, approximately 100 μL of milk were plated onto Columbia CNA agar with 5% sheep blood and MacConkey agar. Agar plates were incubated in aerobic conditions at 37 °C for a total of 42 to 48 h. Plates were examined by a trained technician to evaluate the presence of growth of distinct isolates at 18 to 24 hours and 42 to 48 hours. Samples were defined as contaminated if more than 3 different isolates were identified and omitted from further analysis (Dean et al., 2022). Taxonomic assignments of cultured isolates were made using a Matrix-Assisted Laser Desorption/Ionization-Time of Flight (**MALDI-TOF**) mass spectrometer (**MS**) (Microflex; BrukerDaltonics Inc.) (Jahan et al., 2021). Mass spectra profiles produced from each isolate were matched against the Biotyper reference library. Confidence scores were used to assign genus- and species-level classifications (≥1.8 and ≥ 2.0 for genus- and species-classification, respectively) (Jahan et al., 2021). An IMI was defined as a composite sample containing one or more than one colony forming units (10 CFU/mL) of any cultured isolate.

### Teat apex sample collection

Cows enrolled from two of the participating farms, where labor was available, were sampled weekly from 8 weeks before calving to 5 weeks after calving in the milking parlor prior to the morning milking. First, teats were visually inspected, and any visible debris was removed using a paper towel. Then, using a new pair of disposable gloves, a single sterile pre-moistened gauze was used to thoroughly swab the teat apices of all four teats. After sampling, each gauze was placed in individual tubes containing a pre-loaded solution of 50% glycerol. Gauze samples were transported in a cooler with ice and stored in -80 °C after arrival to research facilities. Postpartum samples were collected before pre-dipping disinfectant solutions were applied to the teat and before milk samples were collected.

### Implementation of case control study

Milk culture results were used to select a subset of gauze samples based on the presence or absence of an IMI caused by SAU or *Streptococcus* spp. and *Streptococcus*-like organisms (**SSLO**) during the first 35 DIM. These mastitis pathogens were used to guide sample selection because they are known to persist in the mammary gland and are strongly associated with elevated SCC and impaired udder health outcomes (Peña-Mosca et al., 2024b). Cows with and without an IMI were individually matched based on farm, lactation number, and the timing of IMI detection (within ±7 DIM). Selected teat apex samples were processed for bacterial culture and species identification as described below. For three cows without a sample available from the week prior to IMI detection, two samples were processed from the closest available time points. In total, 71 samples were collected from 68 cows (34 with an IMI and 34 without an IMI).

### Teat apex sample culture and species identification

The methods for culturing gauze samples collected from teat apex were adapted from protocols utilized in the Laboratory for Udder Health at the University of Minnesota to process towel samples (Rowe et al., 2019). Before sample processing and to reduce the possibility of batch effects, sample IDs for each gauze sample were randomly sorted using Excel (Microsoft Corporation). In accordance with this order, isolates were selected for each sampling processing date in batches of 10 isolates. Gauze samples were thawed at 4 °C overnight and then transferred into 50 mL round-bottom falcon tubes (Corning Inc.). Afterwards, 5 mL of Phosphate-Buffered Saline (**PBS**; Gibco, ThermoFisher Scientific) was added to reduce viscosity. Samples were homogenized carefully by turning tubes upside down twice and vortexing for 10 seconds at low speed. Samples were allowed to stand for 10 minutes, and the homogenization process was repeated once. Following homogenization, a single 1 in 10 serial dilution was done using PBS. A 100 uL aliquot from each sample (i.e. undiluted and 10^-1^) was inoculated onto Mannitol salt agar (Oxoid). Plates were then incubated in aerobic conditions for 24 hours at 37 °C. Following incubation, bacterial groups were visually assessed and counted from the dilution plate with the optimal number of colonies (25 to 250 per plate). Isolates of visually distinct colonies were then picked from the plate using a sterile calibrated loop (Thermo Fisher Scientific), streaked onto blood agar (Hardy Diagnostics) and incubated for 24 hours at 37 °C. After achieving pure growth, the identity of representative colonies was determined using MALDI-TOF MS, using similar methods to those described for isolates identified in milk samples. Non-*aureus staphylococci* and *mammaliicocci* isolates were then preserved at -80 °C using commercial cryovials (Hardy Diagnostics) until further analysis. NASM species-level taxonomy was further investigated through whole genome sequencing, as reported below. Colony forming units (**CFU**) per square inch of gauze were estimated for each distinct colony type based on the number of colonies present on each plate type using a correction factor based on the used dilution. Specifically, a 10x correction factor was applied for 10^-1^ plates, and this calculation also accounted for inoculum size (10x), dilution with PBS (2x), and gauze size (1/4x).

### Determination of in vitro inhibitory activity against Staphylococcus aureus and Streptococcus uberis

The methods utilized in this study were based on procedures previously described by others (Toledo-Silva et al., 2022). For each batch of isolates, on day 1, isolates previously identified as NASM using MALDI-TOF MS were randomly sorted using Excel (Microsoft Corporation). Following this procedure, isolates previously isolated from teat apex were grown for 24 hours in blood agar. If evidence of mixed growth (more than 1 colony type) was present, samples were replated. On day 2, one colony was picked with a 10 uL disposable loop and was grown on brain heart infusion (**BHI**; Research Products International Corporation) media for 18 to 20 hours. The next day, 1 in 10 serial dilutions solutions were created using sterile PBS in a 96 well plate (Thermo Fisher Scientific). Bacterial counts (CFU/mL) were estimated by plating 100 uL of the serial dilutions on blood agar plates that were then aerobically incubated for 18 to 20 h at 37 °C. One hundred microliters of bacterial suspensions containing NASM isolates at different dilutions (10^-2^, 10^-3^, 10^-4^, 10^-5^, 10^-6^, 10^-7^) were added to 10 mL of 2% Tryptic soy agar (**TSA**; Oxoid) inside Falcon round-bottom polypropylene tubes (Corning Inc.). The temperature was examined using a laboratory digital thermometer (Thermco) before the inclusion of the 2% TSA. The media were not used until the temperature reached 50 °C (Toledo-Silva et al., 2022). Tubes were then homogenized and poured into pre-heated (37 °C) TSA plates. Following aerobic incubation for 18 to 20 h, 10 uL of the bacterial suspensions containing approximately 10,000 CFU of SAU (American Type Culture Collection (**ATCC**) strain Seattle 1945; *Staphylococcus aureus* subsp. aureus Rosenbach 25923 ^™^) and *Streptococcus uberis* (**SUB**; ATCC strain 01040 J; *Streptococcus uberis* Diernhofer BAA-854 ^™^) were inoculated onto the top of plates containing different concentrations of NASM isolates and incubated for 18 to 20 h. For each batch of samples, negative controls were included by incubating a 2% TSA agar plate with the inclusion of 100 uL of BHI media only (instead of NASM isolates). Additionally, a 10 μL droplet of PBS was added on top of the media on each plate to serve as an additional negative control, ensuring the absence of PBS contamination. *Staphylococcus aureus* and SUB were incorporated into the media as positive controls, and their self-inhibition was tested to explore the determination of MIC driven by nutrient competition rather than *in vitro* inhibitory activity. Growth of SAU and SUB (i.e., yes vs no) was evaluated by visual examination of each of the plates and comparison with negative control plates within each batch. Minimum inhibitory concentration (**MIC**) was defined as the minimum concentration of NASM (CFU/mL) on the 2% TSA top layer able to inhibit the growth of SAU or SUB.

### DNA extraction, library preparation and whole genome sequencing

Isolates were allowed to grow on BHI for 24 hours, centrifuged at 10,000 rpm for 5 minutes, and the pellet was collected in 2mL sterile polypropylene microcentrifuge tubes (Thermo Fisher Scientific). Following this, DNA extraction was performed utilizing commercial kits (Qiagen’s DNeasy 96 Blood & Tissue Kit, Qiagen). Amount and quality of extracted DNA was evaluated using PicoGreen fluorometer and NanoDrop spectrophotometer (Thermo Fisher Scientific). Library preparation was performed using the DNA Prep Sample Preparation Kit (Illumina Inc.). Resultant DNA was subjected to paired-end sequencing (2x300) utilizing the Illumina MISEQ platform aiming for 50x coverage of *Staphylococcus* spp. genome.

### Bioinformatics

Quality control of raw sequencing data was performed using FastQC (Andrews, 2010) and MultiQC (Ewels et al., 2016). Trimmomatic was utilized to trim low quality base pairs and Nextera XT adapter sequences from raw reads (Bolger et al., 2014). MultiQC was then utilized to summarize the QC metrics during the filtering and trimming process. Trimmed sequence reads were assembled into contigs using SPAdes version 3.15.5 (Bankevich et al., 2012). Assembly quality was evaluated using Quast version 5.2.0 (Gurevich et al., 2013). Taxonomic classification of contigs was performed using GTDB-Tk version 1.7.0 (Chaumeil et al., 2020) and GTDB release 202 (Parks et al., 2022) as implemented in KBase (Arkin et al., 2018). The ABRIcate pipeline (Seemann, 2023a). was used with default settings to identify antimicrobial resistance genes (**ARGs**) from contigs using the MEGARes v2 reference database (Doster et al., 2020). To identify virulence factors from DNA contigs, we used BLAST+ (Camacho et al., 2009) and a custom comprehensive protein database (Naushad et al., 2019), previously shown to be effective for identifying virulence genes in *Staphylococcus* genomes (Fergestad et al., 2021; Naushad et al., 2019). Only BLAST+ results with an e-value of less than or equal to 10 ^e-5^ (Choudhuri, 2014) and amino acid identity of greater than or equal to 35% (Rost, 1999) were considered for analysis. The BAGEL4 online server with default settings was used to identify gene clusters associated with the production of AMPs on contigs (van Heel et al., 2018). Snippy was used with default settings (i.e., minimum coverage of 10x and a minimum variant fraction of 90%) to identify core single nucleotide polymorphisms (Seemann, 2023b) by aligning trimmed reads from each of the isolates onto reference genomes obtained from Genbank (Clark et al., 2016). The following reference genomes were used for each NASM species: *Staphylococcus chromogenes* (**SCH**): GenBank accession number: GCA_002994305.1, *Staphylococcus haemolyticus* (**SHAEM**): GenBank accession number: GCA_001611955.1, *Mammaliicocci sciuri* (**MSCf**): GenBank accession number: GCA_002209165.2, *Staphylococcus succinus* (**SSUC**): GenBank accession number: GCA_001902315.1, *Staphylococcus xylosus* (**SXYL**) and *Staphylococccus pseudoxylosus* (**SPXYL**): GenBank accession number: GCA_000709415.1. As a result of the limited availability of only one or two genomes for *Staphylococcus cohnii* (**SCO**), *Staphylococcus devrieseii* (**SDEV**), and *Staphylococcus equorum* (**SEQ**), SNP calling was not performed on these species. The generated Snippy core files were used to build a maximum likelihood tree using IQ-TREE, with 1,000 bootstrap replicates to assess the robustness of the phylogenetic tree (Nguyen et al., 2015). Trees were visualized and annotated using iTOL (Letunic and Bork, 2021).

### Statistical analysis

All statistical analyses were performed in R version 4.3.2 (https://www.r-project.org/). Code and output are available online: https://fepenamosca.github.io/nasm_antimicrobial_wgs/.

### Association between IMI status and NASM colonization of teat apex

The relationship between IMI status and log_10_ -transformed bacterial counts from teat apex samples was investigated using linear regression models. The relationship between IMI status and presence of NASM on the teat apex samples (i.e., yes vs no) was investigated using a generalized linear model with a “binomial” family and “logit” link (logistic regression). For analysis, NASM taxonomy was determined based on the assigned taxonomy using MALDI-TOF MS and whole genome sequencing. It was represented as a six-level multi-level variable (SCH, SHAEM, MSC, SSUC, SXYL, and other rare NASM [SCO, SDEV, SEQ]).

### Association between IMI status, NASM taxonomy and presence of NASM with in vitro antimicrobial activity on teat apex

Cox proportional hazards regression was used to assess the association between presence of IMI, NASM taxonomy and the concentration to NASM *in vitro* growth inhibition against SAU and SUB. These models were utilized to estimate hazard ratios (**HR**) and their 95% confidence intervals (**CI**), used to compare the inhibitory activity across groups, where higher HR indicates higher NASM inhibitory activity and the presence of inhibition of mastitis pathogens at lower NASM concentrations on the media. As some isolates exhibited no inhibitory activity at the maximum concentration analyzed, observations were right-censored if inhibitory activity remained absent at the highest NASM concentration examined (i.e., 10^-2^ dilution).

Proportional hazards assumption was tested by evaluating the Schoenfeld residuals. Since multiple isolates were obtained from teat apices each cow, the non-independence of observations within each enrolled animal was accounted for using the robust sandwich estimator. The NASM MIC (log_10_, CFU/mL) against SAU and SUB was further explored and compared across cows with and without an IMI and different NASM species using mixed linear regression. These models incorporated cow-ID as a random effect to address the non-independence of observations for isolates obtained from the same cow. Subsequently, isolates were ranked according to their MIC (e.g., 1 to 77) and classified as being in the top 10 isolates with lowest MIC or not (“top 10”). The relationship between IMI status and presence of a “top 10” NASM on the teat apex was investigated using a generalized linear model (logistic regression) with “binomial” family and “logit” link. When performing this analysis by NASM species, we used the Fisher exact test due to the absence of “top 10” isolates for some NASM species. The intraclass correlation coefficient was estimated using the “performance” package (Lüdecke et al., 2021). Estimated marginal means were estimated using the “emmeans” package in R (Lenth et al., 2024) .

Normality assumption was investigated by evaluation of model residuals and the use of quantile-quantile plots. Multiple comparisons were accounted for with Tukey adjustment as implemented in the “emmeans’’ package (Lenth et al., 2024). Farm-ID was forced into all models that investigated the relationship between IMI status and NASM inhibitory activity against SAU and SUB. Because farm ID was strongly associated with the prevalence of different NASM species (i.e., each of the NASM species were in most cases highly prevalent in one farm but rarely found in the other one), this variable was not included in the models comparing *in vitro* inhibitory activity across different NASM species. Processing batch (i.e., the date at which gauze samples were processed or antimicrobial activity of NASM was evaluated) and parity of the cow (i.e., first vs second lactation) were offered as potential confounders. Presence of confounding was evaluated by examining the change in the estimates after adjustment by each confounder ([Crude estimate – Adjusted estimate] / Adjusted estimate). Confounders that changed the estimates by 10% or more were kept in the models.

### Association between the phylogeny, genotypic, and in vitro antimicrobial activity of NASM isolated from the teat apex

Because NASM species were strongly associated with *in vitro* inhibitory activity and presence of AMP gene clusters, all statistical analyses were performed separately for each of the NASM species (SCH, SHAEM, MSC, SSUC, SXYL, SPXYL, Others [SCO, SDEV, SEQ]). A clade, also known as a monophyletic group, was defined as a group of genomes on a phylogenetic tree that included their most recent shared ancestor and their descendants (Kapli et al., 2020). Only AMP gene clusters that were both present and absent in at least 2 isolates within a given species were investigated using regression analysis. For modeling, the association between IMI status, clade membership and the presence of AMP gene clusters was investigated using logistic regression. The presence of AMP gene clusters across different NASM species, and clades within species, was compared using the Fisher exact test. This approach was chosen because, in several cases, gene clusters were found exclusively in a single clade or species. The association between the presence of AMP gene clusters, clade membership, and the MIC of NASM against SAU and SUB for each NASM species was investigated using linear regression.

## RESULTS

### Descriptive characteristics

This study examined two USDA-organic certified farms, one in Minnesota (A) and one in Colorado (B), with varying herd sizes and housing systems. On Farm A, all cows included in this study (14/14) were in their first lactation, while in Farm B, roughly half (43/93) were in their first lactation, and half (50/93) were transitioning to their second lactation. Isolates from 68 cows (34 cows with an IMI and 34 cows without an IMI) from these two farms were included in this study. Fifteen of the cows with an IMI had a SAU-IMI, 15 had a SSLO-IMI, and 4 had an IMI caused by both microorganisms. In the first postpartum composite milk sample, 64.7% (22/34) of the cases were already positive for SAU or SSLO IMI. Teat apex samples from cows with and without an IMI were on average collected on similar days relative to calving (mean ±SD; no IMI: -2.8 ±15.2 vs IMI: -1.2 ±16.5).

### Association between IMI status and NASM colonization of teat apex

NASM were isolated from the teat apex of dairy cows during the transition period in a similar proportion of those without (66.7% [22/36]) and with an IMI (62.9% [24/35]) (OR [95% CI]: 0.85 [0.32, 2.25], *P* = 0.74; **Table 1**). Likewise, NASM counts (log₁₀ CFU/gauze inch²) on the teat apex were similar between groups (Estimate [95% CI]: -0.09 [-0.51, 0.35], *P* = 0.69; **Table 1**). The distribution of NASM species differed by farm. *Staphylococcus chromogenes,* SHAEM, and SSUC showed a high prevalence in farm A (28.1% [16/57], 28.1% [16/57] and 14.1% [8/57], respectively), but were rarely found on farm B (7.1% [1/14], 7.1% [1/14] and 0% [0/14], respectively). On the other hand, SXYL or SPXYL were found in more than half of the samples from farm B (57.1% [8/14]) but showed a low prevalence in farm A (3.5% [2/57]).

**Table 1.** Non-aureus *staphylococci* and *mammaliicocci* counts (log_10_ CFU/square inch of gauze) and prevalence in cows without and with a *Staphylococcus aureus* or *Streptococcus* spp. and *Streptococcus*-like organisms (n=71 teat apex samples).

| Outcome | SAU SSLO IMI |  | Estimate (95%CI) | P-value |
| --- | --- | --- | --- | --- |
|  | No | Yes |  |  |
| Counts |  |  |  |  |
| Overall | 1.10 ± 0.17 | 1.02 ± 0.17 | -0.09 (-0.54, 0.35) | 0.69 |
| *Staph chromogenes* | 0.26 ± 0.11 | 0.20 ± 0.12 | -0.06 (-0.34, 0.23) | 0.71 |
| *Staph haemolyticus* | 0.20 ± 0.13 | 0.21 ± 0.13 | 0.01 (-0.31, 0.32) | 0.96 |
| *Mammaliicoccus sciuri* | 0.19 ± 0.10 | 0.20 ± 0.10 | 0.01 (-0.23, 0.26) | 0.91 |
| *Staph succinus*<sup>a</sup> | 0.17 ± 0.09 | 0.13 ± 0.08 | -0.04 (-0.27, 0.19) | 0.75 |
| *Staph xylosus/pseudoxylosus* | 0.46 ± 0.09 | 0.53 ± 0.10 | 0.07 (-0.17, 0.31) | 0.56 |
| Other | 0.20 ± 0.08 | 0.15 ± 0.08 | -0.05 (-0.24, 0.14) | 0.60 |
| Prevalence |  |  |  |  |
| Overall | 66.7% (24/36) | 62.9% (22/35) | 0.85 (0.32, 2.25) | 0.74 |
| *Staph chromogenes* | 25.0% (9/36) | 22.9% (8/35) | 0.89 (0.29, 2.73) | 0.84 |
| *Staph haemolyticus* | 25.0% (9/36) | 22.9% (8/35) | 0.89 (0.29, 2.73) | 0.84 |
| *Mammaliicoccus sciuri* | 8.3% (3/36) | 5.7% (2/35) | 0.66 (0.08, 4.28) | 0.66 |
| *Staph succinus*<sup>a</sup> | 11.1% (4/36) | 11.4% (4/35) | 1.03 (0.23, 4.72) | 0.97 |
| *Staph xylosus/pseudoxylosus* | 11.1% (4/36) | 17.1% (6/35) | 1.66 (0.43, 7.02) | 0.46 |
| Other<sup>b</sup> | 8.3% (3/36) | 2.9% (1/35) | 0.31 (0.01, 2.63) | 0.29 |

<sup>a</sup> *Staph succinus*: Model did not account for farm, because it was not present in farm B. <sup>b</sup> Other: *Staph cohnii*, *Staph devriesei* and *Staph equorum*. NASM: non-aureus staphylococci and *mammaliicocci*.

### Association between IMI status, NASM taxonomy and presence of NASM with in vitro antimicrobial activity on teat apex

A total of 273 isolates were obtained from the teat apices of the enrolled cows and submitted for identification using MALDI-TOF MS as shown in **Table 2**. Surprisingly, *Bacillus* spp. was the most frequently identified genus. Non-*aureus staphylococci mammaliicocci* were identified in 27.7% (39/141) and 31.8% (42/132) of the isolates obtained from the teat apex samples from cows with and without an IMI, respectively. The most frequent species identified within this group included SCH (8.1% [22/273]) and SHAEM (7.0% [19/273]). Results of MALDI-TOF MS and whole genome sequencing for NASM isolates are presented in supplementary materials (https://doi.org/10.6084/m9.figshare.25832938.v2). All isolates classified as SCH (n=20) or SHAEM (n=15) using MALDI-TOF MS showed similar taxonomy when analyzed using whole genome sequencing. Isolates initially classified as *Staphylococcus xylosus/saprophyticus* (n=12) using MALDI-TOF MS were subsequently reclassified *as* SXYL (n=8) or SPXYL (n=4) by whole genome sequencing. The remaining isolates (34/83; 41.0%) were not identified up to the species-level using MALDI-TOF MS. These isolates were identified using whole genome sequencing as SCH (n=2), SCO (n=1), SDEV (n=2), SEQ (n=1), SHAEM (n=4), SPXYL (n=1), MSC (n=6), SSUC (n=12), and SXYL (n=5).

**Table 2.**
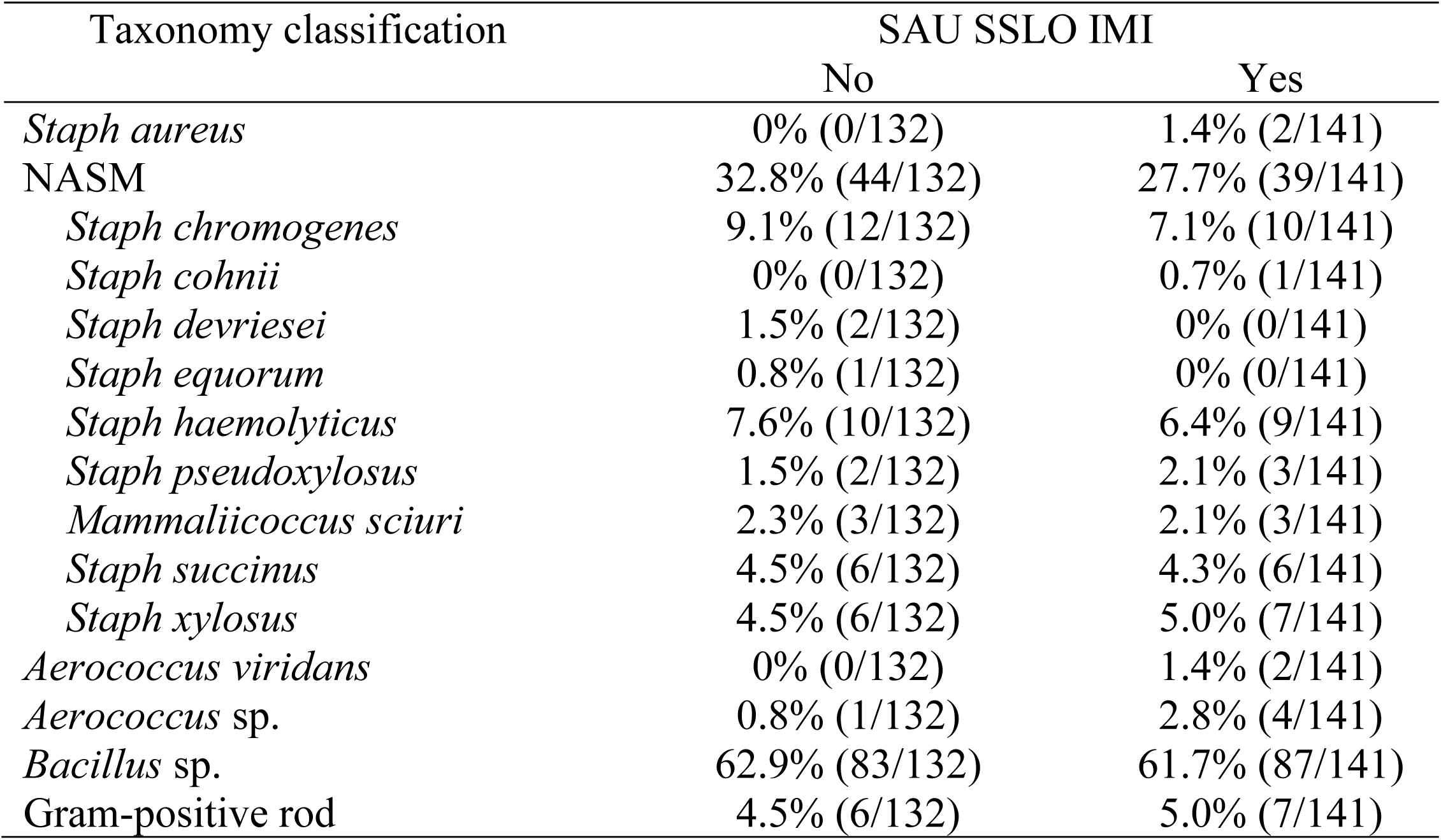
Taxonomic identification of isolates isolated from teat apex samples and submitted for taxonomic identification using Matrix-Assisted Laser Desorption/Ionization-Time of Flight mass spectrometer (n=273 isolates) and confirmation of species-level taxonomy for non-*aureus staphylococci* and *mammaliicocci* (**NASM**) using whole genome sequencing.

We investigated the inhibitory activity of NASM included in a top layer inside the media at varying concentrations against SAU and SUB. In some cases, inhibitory activity was not observed even at the highest concentration tested. In fact, among the 77 NASM isolates evaluated, no inhibitory activity was observed for 5 isolates against SAU and 14 isolates against SUB (**Figure 1**). For these isolates with no inhibitory activity, the highest concentration tested averaged (mean±SD) 3.98±0.45 log₁₀ CFU/mL for SAU and 4.30±0.52 log₁₀ CFU/mL for SUB.

**Figure 1.**
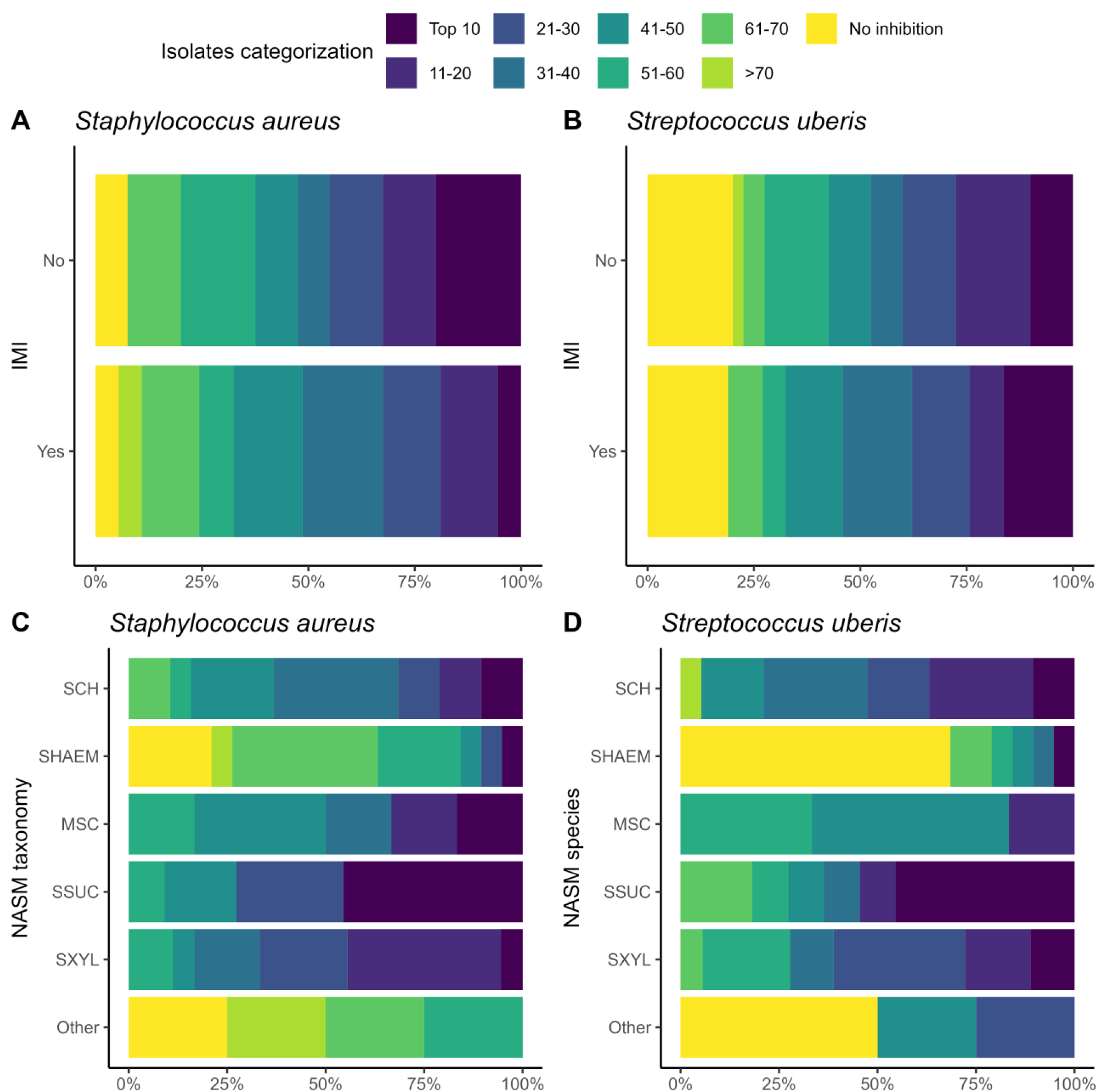
Ranking of 77 non-*aureus staphylococci* and *mammaliicocci* (NASM) isolates by minimum inhibitory concentration (MIC) against *Staphylococcus aureus* and *Streptococcus uberis* isolated from teat apex stratified by presence of intramammary infections by *Staphyloccocus aureus* and *Streptococcus uberis* (SAU SSLO IMI) (panel A/B) and NASM taxonomy, as determined using Matrix-Assisted Laser Desorption/Ionization-Time of Flight and whole genome sequencing (panel C/D) (n=79 isolates). No inhibitory activity: NASM isolates that showed growth of *Staphylococcus aureus* or *Streptococcus uberis* at the maximum investigated concentration on media. SCH: *Staphylococcus chromogenes*, SHAEM: *Staphylococcus haemolyticus*, MSC: *Mammaliicoccus sciuri*, SSUC: *Staphylococcus succinus,* SXYL: *Staphylococcus xylosus/pseudoxylosus,* Other: *Staphylococcus cohnii, Staphylococcus devriesei* and *Staphylococcus equorum*.

For isolates where inhibition was observed, MIC values did not differ between those from cows with and without an IMI (**Table 3**). To account for the varying concentrations required to reach inhibition and the presence of right-censored data (i.e., isolates without inhibition at the maximum NASM concentration tested), we analyzed the data using survival analysis. The overall presence of inhibitory activity was not different between isolates from cows with and without an IMI (HR [95%CI] for SAU: 0.89 [0.56, 1.45]; for SUB: 1.17 [0.68, 1.97]; **Figure 2**).

**Figure 2.**
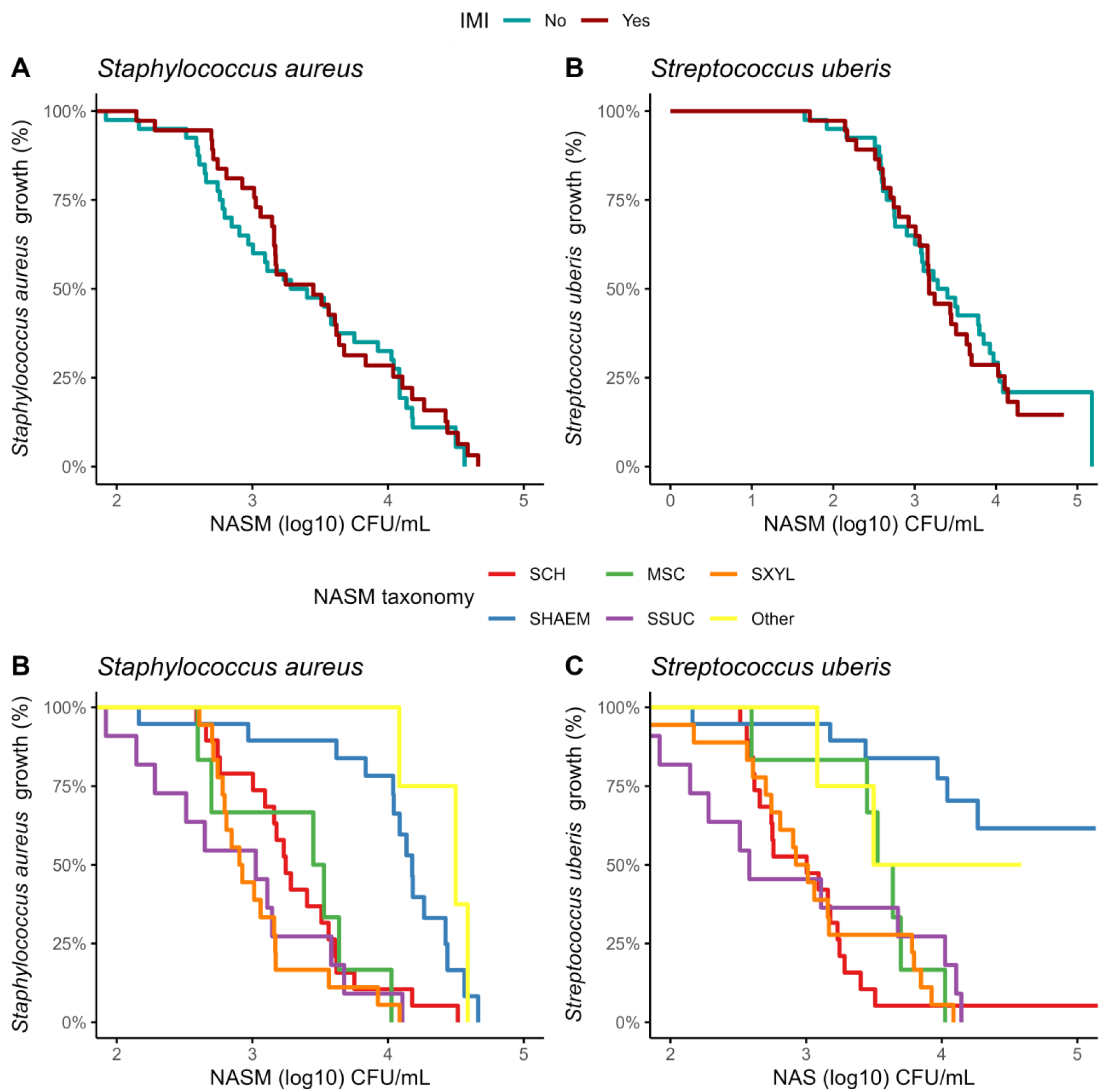
Kaplan-Meier plots showing the presence of *Staphylococcus aureus* and *Streptococcus uberis* growth on top of the media at different concentrations of non-*aureus staphylococci* and mammaliicocci (NASM) in the top layer, stratified by the presence of intramammary infections caused by *Staphylococcus aureus* and *Streptococcus uberis* and NASM taxonomy, as determined using Matrix-Assisted Laser Desorption/Ionization-Time of Flight and whole genome sequencing (n=77 isolates). SCH: *Staphylococcus chromogenes*, SHAEM: *Staphylococcus haemolyticus*, MSC: *Mammaliicoccus sciuri*, SSUC: *Staphylococcus succinus,* SXYL: *Staphylococcus xylosus/pseudoxylosus,* Other: *Staphylococcus cohnii, Staphylococcus devriesei* and *Staphylococcus equorum*.

**Table 3.**
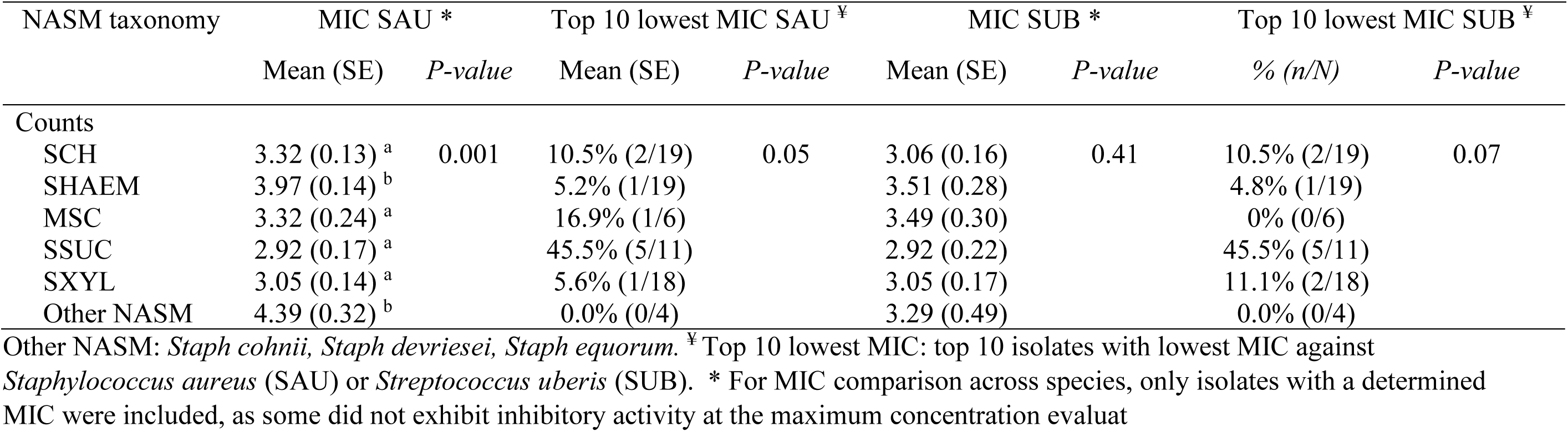
Relationship between species identification and minimum inhibitory concentration (MIC; log CFU/mL) of non-*aureus staphylococci* and *mammaliicocci* (NASM) isolated from teat apex against *Staphylococcus aureus* and *Streptococcus uberis* (n=77 isolates).

| NASM taxonomy | MIC SAU * |  | Top 10 lowest MIC SAU ‡ |  | MIC SUB * |  | Top 10 lowest MIC SUB ‡ |  |
| --- | --- | --- | --- | --- | --- | --- | --- | --- |
|  | Mean (SE) | P-value | Mean (SE) | P-value | Mean (SE) | P-value | % (n/N) | P-value |
| Counts |  |  |  |  |  |  |  |  |
| SCH | 3.32 (0.13) <sup>a</sup> | 0.001 | 10.5% (2/19) | 0.05 | 3.06 (0.16) | 0.41 | 10.5% (2/19) | 0.07 |
| SHAEM | 3.97 (0.14) <sup>b</sup> |  | 5.2% (1/19) |  | 3.51 (0.28) |  | 4.8% (1/19) |  |
| MSC | 3.32 (0.24) <sup>a</sup> |  | 16.9% (1/6) |  | 3.49 (0.30) |  | 0% (0/6) |  |
| SSUC | 2.92 (0.17) <sup>a</sup> |  | 45.5% (5/11) |  | 2.92 (0.22) |  | 45.5% (5/11) |  |
| SXYL | 3.05 (0.14) <sup>a</sup> |  | 5.6% (1/18) |  | 3.05 (0.17) |  | 11.1% (2/18) |  |
| Other NASM | 4.39 (0.32) <sup>b</sup> |  | 0.0% (0/4) |  | 3.29 (0.49) |  | 0.0% (0/4) |  |

Other NASM: *Staph cohnii*, *Staph devriesei*, *Staph equorum*. ‡ Top 10 lowest MIC: top 10 isolates with lowest MIC against *Staphylococcus aureus* (SAU) or *Streptococcus uberis* (SUB). \* For MIC comparison across species, only isolates with a determined MIC were included, as some did not exhibit inhibitory activity at the maximum concentration evaluated

To provide a reference for interpreting NASM’s inhibitory activity, we tested SAU and SUB against themselves by plating each pathogen on media containing varying concentrations of the same microorganism. When SAU was tested as a control on the 2% TSA top layer, its MIC was 3.42 log₁₀ CFU/mL, and 51.9% (40/77) of NASM isolates had an MIC below this value, including 52.5% (21/40) of isolates from cows without an IMI and 51.4% (19/37) from cows with an IMI. Similarly, when SUB was evaluated, its MIC was 2.91 log₁₀ CFU/mL, and 32.4% (25/77) of NASM isolates had an MIC below this threshold. Among these isolates, 35.0% (14/40) were collected from cows without an IMI, and 29.7% (11/37) were from cows with an IMI Building on these MIC distributions, we ranked the isolates based on their inhibitory activity against SAU and SUB to identify those with the highest inhibitory activity (**Figure 1**; **Table 3**). The “top 10” isolates with the lowest MIC (log₁₀ CFU/mL) had a mean ± SD MIC of 2.41 ± 0.26 for SAU and 2.16 ± 0.32 for SUB, with MICs ranging from 1.92 to 2.66 for SAU and 1.65 to 2.56 for SUB. These highly inhibitory isolates were more common in cows without an IMI (19.4% [7/36]) compared to those with an IMI (5.8% [2/35]) (OR [95% CI]: 0.25 [0.04, 1.13], *P* = 0.07). However, an association with IMI status was not observed for cows harboring a “top 10” isolate for SUB (controls: 11.1% [4/36] vs. cases: 14.3% [5/35]; OR [95% CI]: 1.33 [0.32, 5.83], *P* = 0.69).

The inhibitory activity of NASM against SAU and SUB varied significantly among NASM species (*P* < 0.001; **Figure 2**). *Staphylococcus haemolyticus* and other less common NASM species (e.g., SCO, SDEV, SEQ) exhibited lower inhibitory activity against SAU compared to SCH (HR [95% CI]: 0.29 [0.12, 0.59], *P* = 0.02; and 0.15 [0.03, 0.71], *P* < 0.01, *respectively*), MSC (0.21 [0.07, 0.58], *P* <0.001; and 0.12 [0.03, 0.52], *P* < 0.001, respectively), SSUC (0.15 [0.05, 0.42], *P* < 0.001; and 0.08 [0.02, 0.38], *P* < 0.001, respectively), and SXYL/SPXYL (0.13 [0.05, 0.38], *P* < 0.001; and 0.08 [0.02, 0.29], *P* < 0.001, respectively). The adjusted MIC (log₁₀ CFU/mL) of NASM against SAU across species ranged from 2.92 to 4.39 (**Table 3**).

Similarly, when inhibition of SUB was assessed, SHAEM displayed reduced inhibitory activity compared to SCH (HR [95% CI]: 0.10 [0.02, 0.50]), MSC (0.17 [0.06, 0.54]), SSUC (HR [95% CI]: 0.13 [0.03, 0.55]), and SXYL/SPXYL (HR [95% CI]: 0.11 [0.03, 0.47], all *P* < 0.001). The inhibitory activity against SUB of rare NASM species was not different from that of other NASM species identified in this study (**Figure 2**). The adjusted MIC (log₁₀ CFU/mL) of NASM species against SUB ranged between 2.92 and 3.51 (**Table 3**).

The distribution of isolates ranked by MIC against SAU and SUB is shown in **Figure 3**. The proportion of "top 10" inhibitory isolates differed across NASM species (*P* = 0.05 for SAU and *P* = 0.07 for SUB; **Table 3**). Among the species with lower inhibitory activity, only 5.3% (1/19) of SHAEM isolates and 0% (0/5) of rare NASM species isolates were classified as "top 10" against SAU and/or SUB. In contrast, 45.5% (5/11) of SSUC isolates were “top 10” inhibitory isolates against both SAU and SUB (**Figure 3** and **Table 3**)

**Figure 3.**
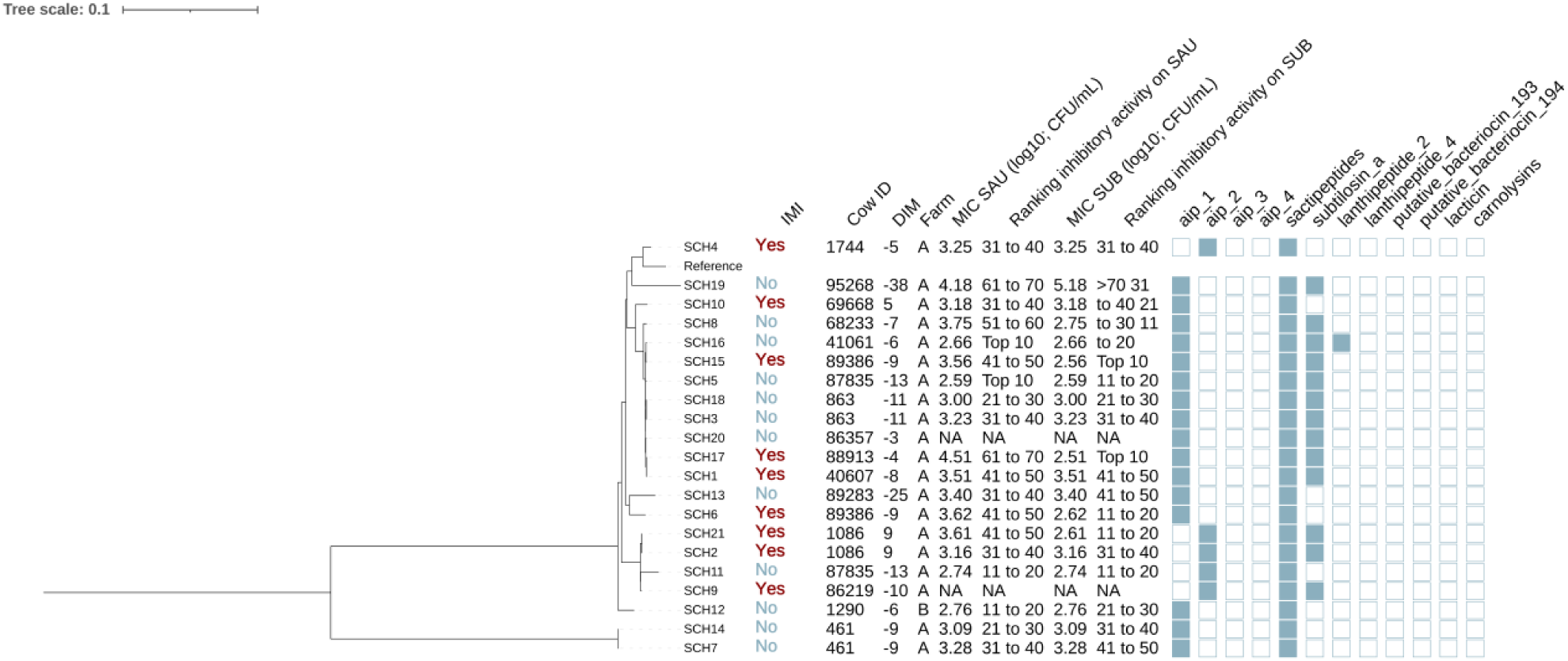
Maximum likelihood phylogenetic tree including 21 isolates identified as *Staphylococcus chromogenes* (SCH). This tree represents a total of 64,794 single nucleotide polymorphisms. Reference: GenBank accession number: GCA_002994305.1. MIC SAU: Minimum inhibitory concentration against *Staphylococcus aureus*. MIC SUB: Minimum inhibitory concentration against *Streptococcus uberis*. Ranking inhibitory activity on SAU: Ranking of isolates according to MIC on SAU. Ranking inhibitory activity on SUB: Ranking of isolates according to MIC on SUB. Auto induced peptide class I (aip_1): NCBI accession number: WP_001093929.1. Auto induced peptide class II (aip_2). NCBI accession number WP_001094921.1. Auto induced peptide class III (aip_3): CBI accession number WP_000735197.1. Auto induced peptide class IV (aip_4): NCBI accession number: WP_001094303.1). Sactipeptides: Interpro accession number PF04055. Subtilosin A NCBI accession number: NP_391616.1. Lanthipeptides class II Interpro accession number: PF05147. Lanthipeptides class IV (Uniprot accession number: O88037). putative bacteriocins 193.2 and 194.2 (putative_bacteriocin_193/194): Genbank accession number AJ002203.2. lacticin Z (lacticin): Interpro accession number PF11758. Carnolysins: Interpo accession number PF00082.

### Association between the phylogeny, genotypic, and in vitro antimicrobial activity of NASM isolated from the teat apex

#### Assembly quality of NASM isolates submitted for WGS

All isolates classified as NASM were submitted for WGS. After quality filtering the genomes were assembled using spades. Across the NASM assembled genomes, an average ± SD of 48.4 ± 137 contigs were identified. The mean ± SD genome size was 2,610,000 ± 276,000, with the N50 value averaging 585,000 ± 485,000 across all genomes. The longest contig exhibited an average size of 883,000 ± 367,000 across all genomes.

#### Phylogeny and antimicrobial activity

Five phylogenetic trees were constructed representing six different NASM species (SCH, SHAEM, MSC, SSUC, SXYL/SPXYL). The assigned NASM taxonomy showed a strong association with the presence of various AMP gene clusters. For example, it was observed that AIP class I gene clusters (NCBI accession number: WP_001093929.1) were present in all isolates identified as SCH and SHAEM, as well as in half of those identified as MSC, but were absent in SSUC, PXYL, and SPSXYL (*P*<0.001). Similarly, putative bacteriocin 193.2 (Genbank accession number: AJ002203.2) gene clusters were highly prevalent in SXYL or SEQ, but were not detected in other NASM species (*P*<0.001).

#### Staphylococcus chromogenes

Among the 21 SCH isolates (**Figure 3**), 12 and 9 were from cows without and with IMI, respectively. The majority (95.2% [20/21]) were from farm A. Genes for AIP class I and II (NCBI accession: WP_001094921.1) were detected in 76.2% (16/21) and 23.8% (5/21) of SCH isolates, respectively. Sactipeptides (InterPro: PF04055) and subtilosin A (NCBI: NP_391616.1) gene clusters were found in 100% (21/21) and 61.9% (13/21) of SCH isolates. Lanthipeptides class II (Interpro accession number: PF05147) gene clusters were identified in 1 out of the 21 isolates (4.8%). AIP class I and II genes (NCBI: WP_001094921.1) were detected in 76.2% [16/21] and 23.8% [5/21] of isolates, respectively. AIP class I was more frequent in isolates from cows without IMI (91.7% [11/12]) than with IMI (55.6% [5/9]) (*P* = 0.08), while an opposite pattern was observed for AIP class II (no IMI: 8.3% [1/12] vs. IMI: 44.4% [4/9]; *P* = 0.08).

Sactipeptides (InterPro accession number: PF04055) were present in all isolates. Subtilosin A (NCBI: NP_391616.1) was detected in isolates from cows without IMI (41.7% [5/12]) and with IMI (33.3% [3/9]) (*P* = 0.70). Lanthipeptides class II (InterPro: PF05147) were rare (4.8% [1/21]). None of these gene clusters were associated with MIC against SAU or SUB (P > 0.05 for all linear regression models).

Phylogenetic analysis of the 21 SCH genomes revealed two major clades, with one clade containing just two isolates, both from the same cow (SCH7 and SCH14 from cow 46, **Figure 3**). The largest subclade contained 9 isolates (SCH1, SCH3, SCH5, SCH8, SCH15, SCH16, SCH17, SCH18, SCH20). The genomes within this clade were characterized by the consistent presence of subtilosin A (100% [9/9]) and AIP class I (100% [9/9]) gene clusters, which were less common among other SCH isolates (subtilosin A: 33% [4/12], *P* < 0.001; AIP class I: 58.3% [7/12], *P* = 0.045). In contrast, AIP class II gene clusters were not identified in this clade but identified in 41.7% (5/12) of others (*P* = 0.045). This clade also contained the only 2 SCH isolates classified as “top 10” with highest antimicrobial activity. However, MIC (CFU/mL) against SAU or SUB did not differ between this clade and others (SAU: estimate [95%CI]: 0.05 [-0.45, 0.56]), *P*=0.82; SUB: -0.35 [-0.93, 0.22], *P*=0.51). These 21 isolates were obtained from 16 cows, with 5 cows carrying two SCH isolates each. In some cases, both isolates from the same cow showed minimal divergence and clustered within the same clade (e.g., cow 461: SCH7 and SCH14; cow 863: SCH3 and SCH18; cow 1086: SCH2 and SCH21). In others, isolates from different clades were recovered from the same cow on the same day (e.g., cow 87835: SCH5 and SCH11; cow 89386: SCH5 and SCH15), suggesting the presence of distinct strains within the SCH population colonizing the teat apex.

#### Staphylococcus haemolyticus

A total of 19 isolates were identified as SHAEM (**Figure 4**). Out of these isolates, 10 and 9, were from cows without and with an IMI, respectively. Most, 94.7% [18/19]) were from farm A. Genes encoding AIP class I and sactipeptides were found in all isolates (100% [19/19]), while lanthipeptides class IV were detected in 5.3% [1/19]

**Figure 4.**
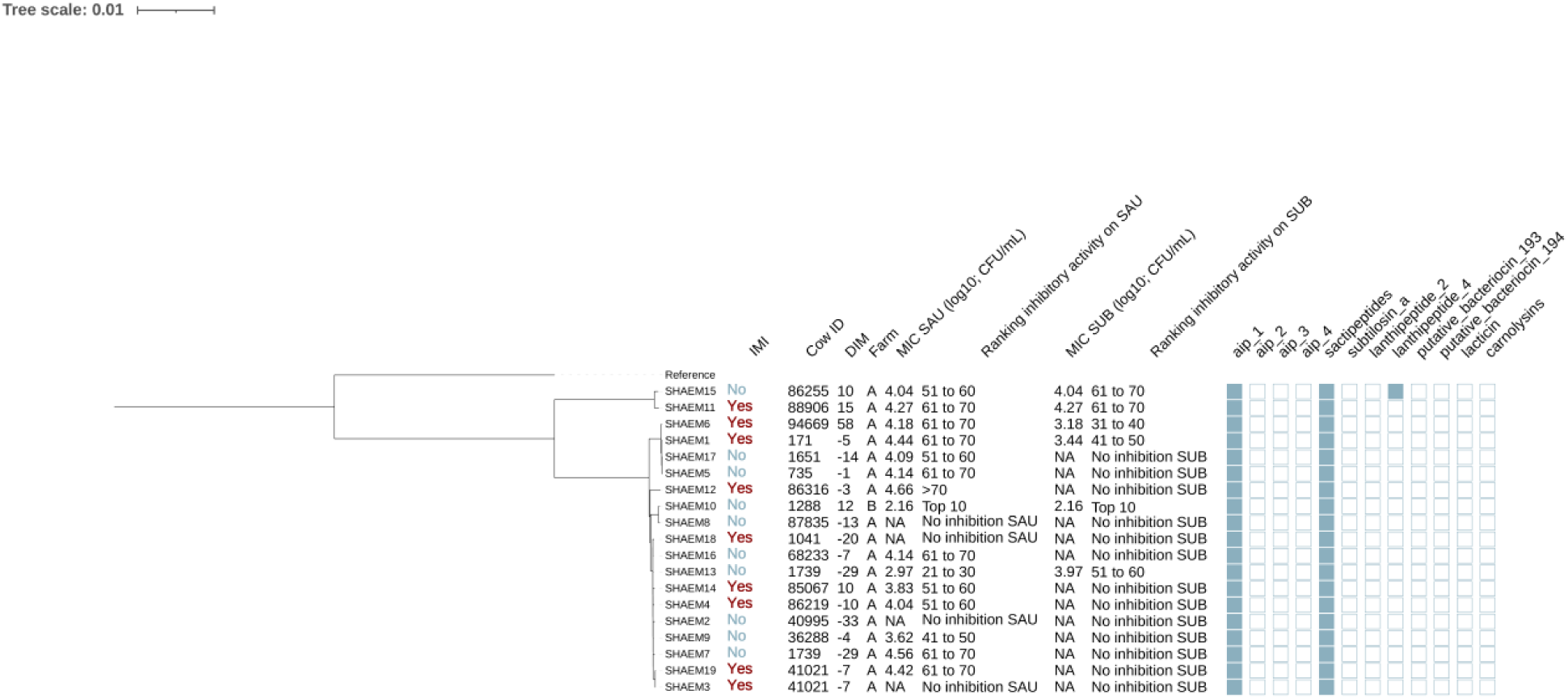
Maximum likelihood phylogenetic tree including 19 isolates identified as *Staphylococcus haemolyticus* (SHAEM). This tree represents a total of 52,188 single nucleotide polymorphisms. Reference: GenBank accession number: GCA_001611955.1. MIC SAU: Minimum inhibitory concentration against *Staphylococcus aureus*. MIC SUB: Minimum inhibitory concentration against *Streptococcus uberis*. Ranking inhibitory activity on SAU: Ranking of isolates according to MIC on SAU. Ranking inhibitory activity on SUB: Ranking of isolates according to MIC on SUB. Auto induced peptide class I (aip_1): NCBI accession number: WP_001093929.1. Auto induced peptide class II (aip_2). NCBI accession number WP_001094921.1. Auto induced peptide class III (aip_3): CBI accession number WP_000735197.1. Auto induced peptide class IV (aip_4): NCBI accession number: WP_001094303.1). Sactipeptides: Interpro accession number PF04055. Subtilosin A NCBI accession number: NP_391616.1. Lanthipeptides class II Interpro accession number: PF05147. Lanthipeptides class IV (Uniprot accession number: O88037). putative bacteriocins 193.2 and 194.2 (putative_bacteriocin_193/194): Genbank accession number AJ002203.2. lacticin Z (lacticin): Interpro accession number PF11758. Carnolysins: Interpo accession number PF00082.

The 19 SHAEM isolates were grouped into 2 distinct clades, with one clade comprising just two isolates, as shown in **Figure 4**. The largest subclade contained 10 isolates: SHAEM2, SHAEM3, SHAEM4, SHAEM7, SHAEM9, SHAEM13, SHAEM14, SHAEM16, SHAEM18, SHAEM19. Half of these isolates were obtained from cows without IMI (5/10) and half from cows with an IMI (5/10). Like other SHAEM isolates, this clade showed limited inhibitory activity, with many unable to inhibit SAU (30.0% [3/10]) or SUB (90.0% [9/10]). Interestingly, a small subclade containing 2 isolates (SHAEM 8 and SHAEM10), encompassed isolates obtained from cows in different farms and exhibiting different inhibitory activity against SAU or SUB. We did not investigate potential associations with genes associated with AMP production because they either had low prevalence or were present in all SHAEM isolates.

#### Mammaliicoccus sciuri

Six isolates were identified as MSC (**Figure 5**). Half of these isolates were cultured from controls (3/6) and half from cases (3/6). In addition, 4 of these isolates were obtained from cows in farm A and 2 from cows in farm B. The genomes of these isolates contained genes that encoded AIPs class I (no IMI: 66.7% [2/3], IMI: 33.3% [1/3]) (*P*=0.42), sactipeptides (present in all isolates) and putative bacteriocin 193.2 (Genbank accession number: AJ002203.2) (no IMI: 33.3% [1/3], IMI: 66.7% [2/3], P=1.00). No associations were found between presence of these genes, and the MIC against SAU or SUB (*P*>0.05 for all linear models).

**Figure 5.**
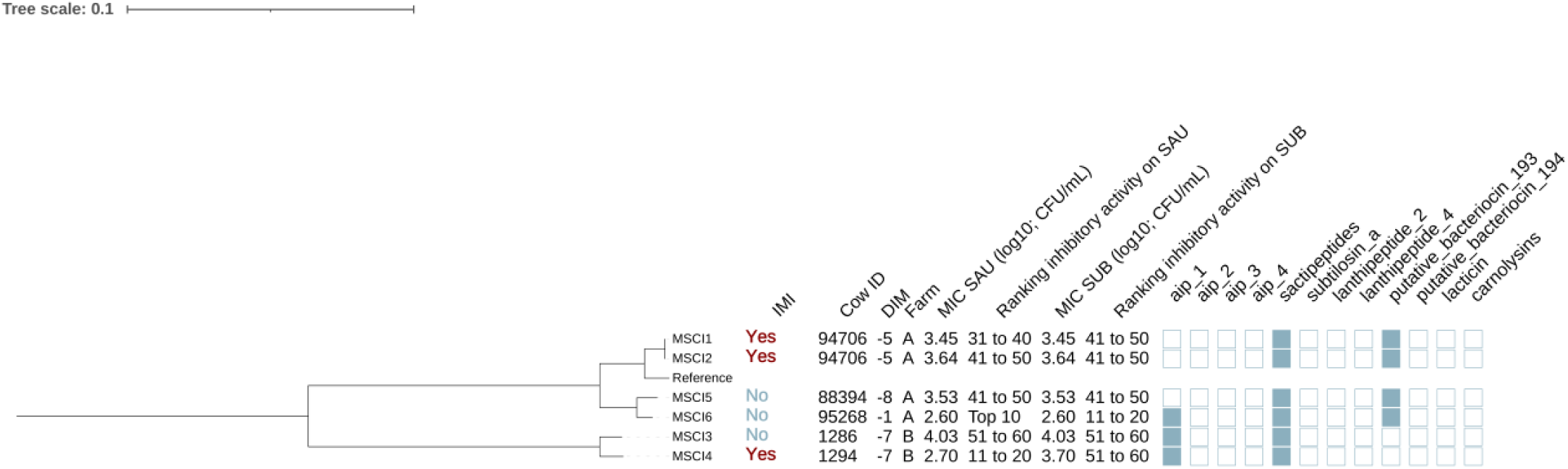
Maximum likelihood phylogenetic tree including 6 isolates identified as *Mammaliicoccus sciuri* (MSC). This tree represents a total of 93,312 single nucleotide polymorphisms. Reference: GenBank accession number: GCA_002209165.2. MIC SAU: Minimum inhibitory concentration against *Staphylococcus aureus*. MIC SUB: Minimum inhibitory concentration against *Streptococcus uberis*. Ranking inhibitory activity on SAU: Ranking of isolates according to MIC on SAU. Ranking inhibitory activity on SUB: Ranking of isolates according to MIC on SUB. Auto induced peptide class I (aip_1): NCBI accession number: WP_001093929.1. Auto induced peptide class II (aip_2). NCBI accession number WP_001094921.1. Auto induced peptide class III (aip_3): CBI accession number WP_000735197.1. Auto induced peptide class IV (aip_4): NCBI accession number: WP_001094303.1). Sactipeptides: Interpro accession number PF04055. Subtilosin A NCBI accession number: NP_391616.1. Lanthipeptides class II Interpro accession number: PF05147. Lanthipeptides class IV (Uniprot accession number: O88037). putative bacteriocins 193.2 and 194.2 (putative_bacteriocin_193/194): Genbank accession number AJ002203.2. lacticin Z (lacticin): Interpro accession number PF11758. Carnolysins: Interpo accession number PF00082.

Phylogenetic analysis showed that isolates were grouped in 2 distinct clades, one including isolates from farm A and the other one isolates from farm B (**Figure 5**). Isolates from farm A formed two different subclades, one including two isolates obtained from cows with IMI (MSC1 and MSC2) and the other from cows without an IMI (MSCI5 and MSCI6). Putative bacteriocin 193.2 was identified in all isolates isolated from cows in farm A (4/4) but not identified in those from farm B (0/2) (*P*=0.08). No differences were found in the MIC against SAU or SUB across these clades (*P*=0.92 and *P*=0.21, respectively).

#### Staphylococcus succinus

A total of 12 isolates were identified as SSUC (**Figure 6**). Half (6/12) were isolated from cows without IMI, and the remaining half (6/12) from cows with IMI. Furthermore, all SSUC isolates were found in cows from farm A. Genes encoding AIPs class II and Sactipeptides were identified in 100% (19/19) of the isolates. Carnolysins were identified in 1 out of the 12 isolates (8.3%). *Staphylococcus succinus* isolates were grouped in multiple clades (**Figure 4**). One of these clades contained isolates SSUC1, SSUC2, SSUC3, SSUC4, SSUC5, SSUC10 and SSUC12, and the MICs of these isolates were lower than SSUC isolates in other clades (estimate [95%CI]: MIC SAU: -1.06 [-1.66, -0.46], *P*=0.003; MIC SUB: -1.06 [-2.14, 0.02], *P*=0.05). This clade also contained all of the SSUC isolates classified as “top 10” with lowest MIC against both SAU and SUB. Lastly, an interesting finding was that there were 2 cows from which multiple isolates were recovered (cow 94706: SSUC1, SSUC7 and SSUC12; cow 1333: SSUC6 and SSUC10), and these isolates were distributed across distinct subclades, indicating the presence of distinct strains within the teat apex of a given cow. Associations between the presence of genes related to the production of AMPs were not investigated because they showed a low prevalence or were present in all SSUC isolates.

**Figure 6.**
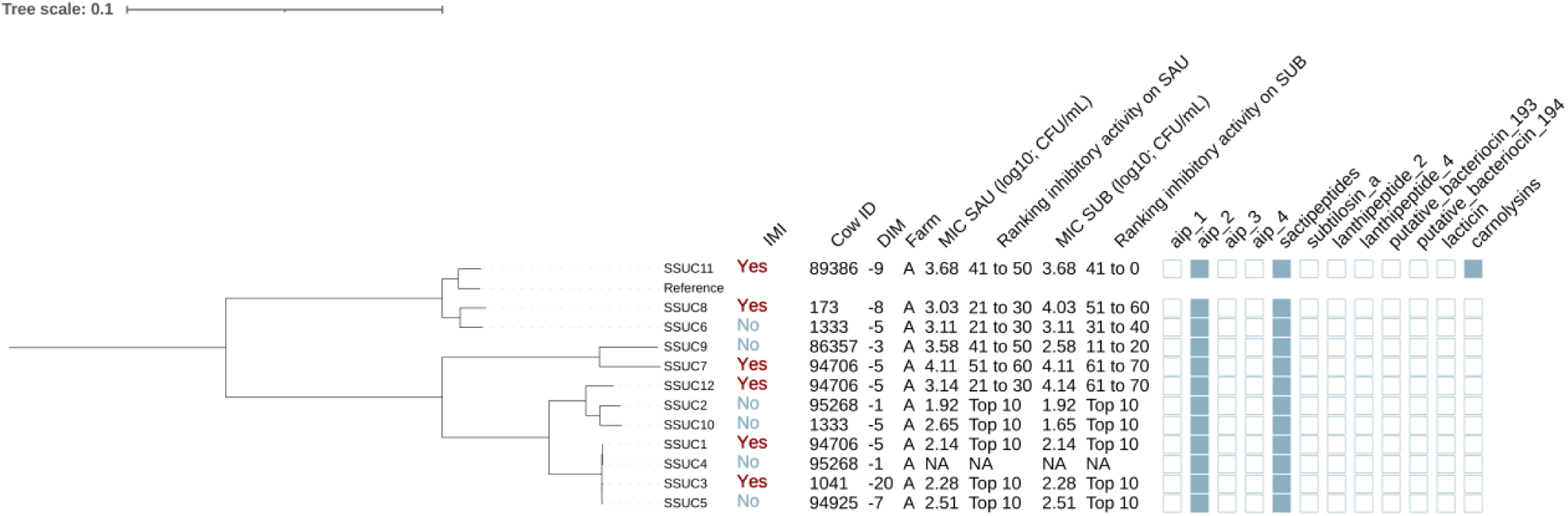
Maximum likelihood phylogenetic tree including 12 isolates identified as *Staphylococcus succinus* (SSUC). This tree represents a total of 69,716 single nucleotide polymorphisms. Reference: GenBank accession number: GCA_001902315.1. MIC SAU: Minimum inhibitory concentration against *Staphylococcus aureus*. MIC SUB: Minimum inhibitory concentration against *Streptococcus uberis*. Ranking inhibitory activity on SAU: Ranking of isolates according to MIC on SAU. Ranking inhibitory activity on SUB: Ranking of isolates according to MIC on SUB. Auto induced peptide class I (aip_1): NCBI accession number: WP_001093929.1. Auto induced peptide class II (aip_2). NCBI accession number WP_001094921.1. Auto induced peptide class III (aip_3): CBI accession number WP_000735197.1. Auto induced peptide class IV (aip_4): NCBI accession number: WP_001094303.1). Sactipeptides: Interpro accession number PF04055. Subtilosin A NCBI accession number: NP_391616.1. Lanthipeptides class II Interpro accession number: PF05147. Lanthipeptides class IV (Uniprot accession number: O88037). putative bacteriocins 193.2 and 194.2 (putative_bacteriocin_193/194): Genbank accession number AJ002203.2. lacticin Z (lacticin): Interpro accession number PF11758. Carnolysins: Interpo accession number PF00082.

#### Staphylococcus xylosus & Staphylococcus pseudoxylosus

We identified 13 isolates as SXYL and 5 as SPXYL (**Figure 7**). Among SXYL isolates, 53.8% [7/13] were collected from cows without IMI and 46.2% [6/13] from cows with IMI. For SPXYL, 60.0% [3/5] were harbored from cows without IMI and 40.0% [2/5] with IMI. Most SXYL (92.3% [12/13]) and SPXYL (80.0% [4/5]) isolates were from farm B. The phylogenetic tree showed that isolates could be divided into 3 distinct clades, two of which contained isolates from SXYL (SXYL clade A: SXYL1, SXYL2, SXYL4, SXYL10, SXYL11, SXYL13, SXYL14; SXYL clade B: other SXYL isolates) and the other of which contained isolates classified as PSXYL. The MIC (log_10_ CFU/mL) against SAU was (mean±SE) 3.14±0.18, 2.98±0.17, and 3.06±0.20 for isolates in SXYL clade A and B and for those identified as SPXYL, respectively (Type III *P-value* = 0.81). In addition, for isolates belonging to SXYL clades A and B and those identified as SPXYL, the MIC against SUB was 3.12±0.26, 2.97±0.28, and 3.06±0.30, respectively (Type III *P-value* = 0.92). Sactipeptides and subtilosin A were present in all SXYL and SPXYL genomes. On the other hand, AIPs class II were present in 53.8% (7/13) of the SXYL isolates (clade A: 33.3% [2/6]; clade B: 71.4% [5/7] and 100% [5/5] of the PSXYL isolates) (*P*=0.07). In addition, AIPs class III (NCBI accession number: WP_000735197.1) were highly prevalent in SXYL clade A (66.7% [4/6]); showed a low prevalence in SXYL clade B (14.3% [1/7]); and were not identified in SPXYL genomes (*P*=0.03). Putative bacteriocin 193.2 genes were identified in all isolates from SXYL clade B (100%, 6/6), had a lower prevalence in SXYL clade A (14.3%, 1/7), and were not found in SPXYL isolates (*P*<0.001). None of these genes were associated with MIC against SAU or SUB (*P*>0.05 for all linear regression models).

**Figure 7.**
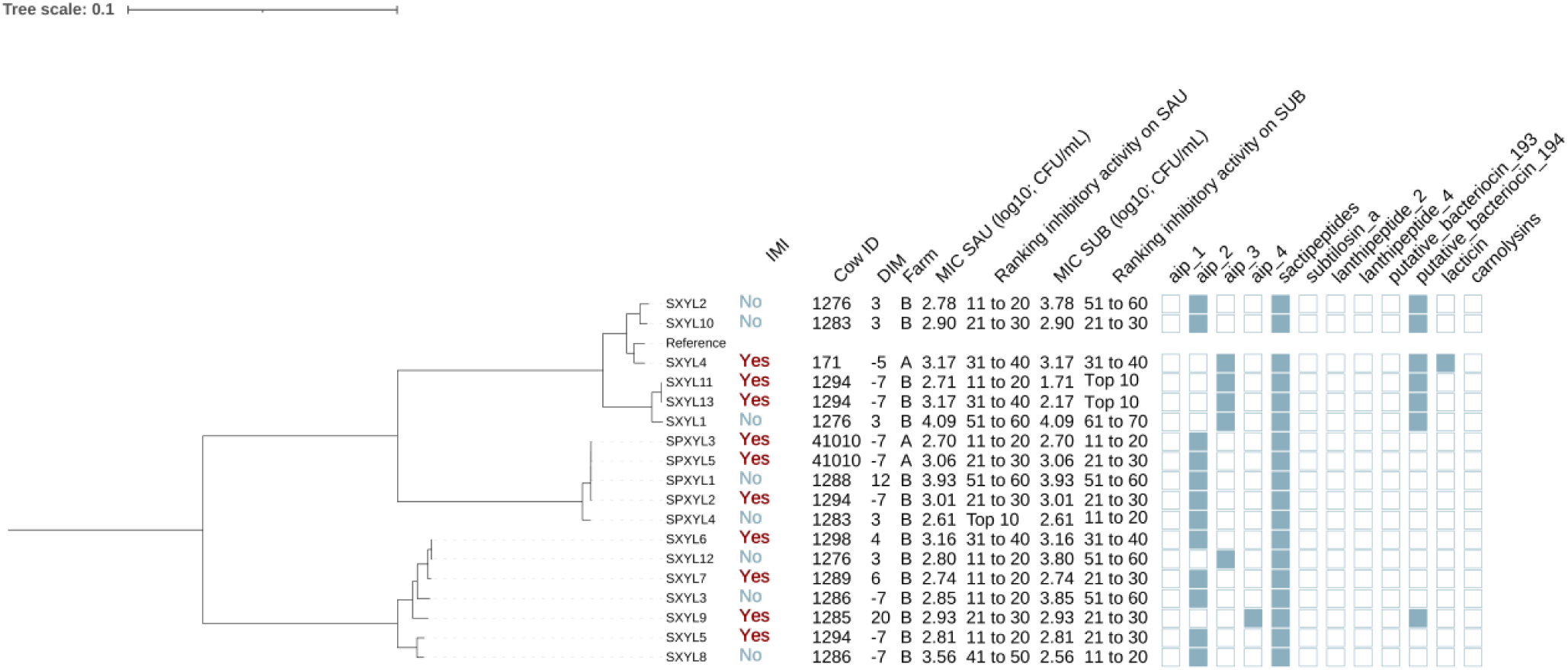
Maximum likelihood phylogenetic tree including 18 isolates identified as *Staphylococcus xylosus* (XYL) and *Staphylococcus pseudoxylosus* (PXYL). This tree represents a total of 209,336 single nucleotide polymorphisms. Reference: GenBank accession number: GCA_000709415.1. MIC SAU: Minimum inhibitory concentration against *Staphylococcus aureus*. MIC SUB: Minimum inhibitory concentration against *Streptococcus uberis*. Ranking inhibitory activity on SAU: Ranking of isolates according to MIC on SAU. Ranking inhibitory activity on SUB: Ranking of isolates according to MIC on SUB. Auto induced peptide class I (aip_1): NCBI accession number: WP_001093929.1. Auto induced peptide class II (aip_2). NCBI accession number WP_001094921.1. Auto induced peptide class III (aip_3): CBI accession number WP_000735197.1. Auto induced peptide class IV (aip_4): NCBI accession number: WP_001094303.1). Sactipeptides: Interpro accession number PF04055. Subtilosin A NCBI accession number: NP_391616.1. Lanthipeptides class II Interpro accession number: PF05147. Lanthipeptides class IV (Uniprot accession number: O88037). putative bacteriocins 193.2 and 194.2 (putative_bacteriocin_193/194): Genbank accession number AJ002203.2. lacticin Z (lacticin): Interpro accession number PF11758. Carnolysins: Interpo accession number PF00082.

#### Genetic determinants of virulence

We investigated the prevalence of virulence genes associated with adherence, exoenzymes, immune evasion, iron uptake, and toxin production across NASM and SAU, with SAU carrying a higher number of these genes. Among adherence-related genes (n=37), 81.1% were detected, with most highly prevalent in SAU, while NASM species showed lower prevalence but retained some key adhesion and biofilm formation genes (**Figure 8**). Of the 22 exoenzyme genes examined, SAU carried 77.3%, whereas NASM species harbored fewer, though nucleases and aureolysins were widely present (**Figure 8**). Immune evasion genes (n=35) were nearly ubiquitous in SAU (97.1%) but varied across NASM, with capsular genes widespread and immune evasion factors such as staphylococcal binding immunoglobulin protein and staphylococcal complement inhibitor mainly found in SAU and SCH (**Figure 9**). For iron uptake and metabolism (n=29), all genes were present in SAU, while NASM species harbored fewer, with prevalence varying distribution (**Figure 9**). Among 87 enterotoxin and exotoxin genes, 59 were identified, with enterotoxins exclusive to SAU and exotoxins detected in SAU and SCH but absent in other NASM (**Figure 10**). Of the 36 toxin-related genes, SAU carried 75.0%, while NASM species had fewer, except for some cases such as exfoliative toxin type C, which was present in all NASM isolates (**Figure 10**), beta phenol-soluble modulins, which were widespread except in SCH and MSC, and type VII secretion system proteins, primarily detected in SCH (**Figure 10**).

**Figure 8.**
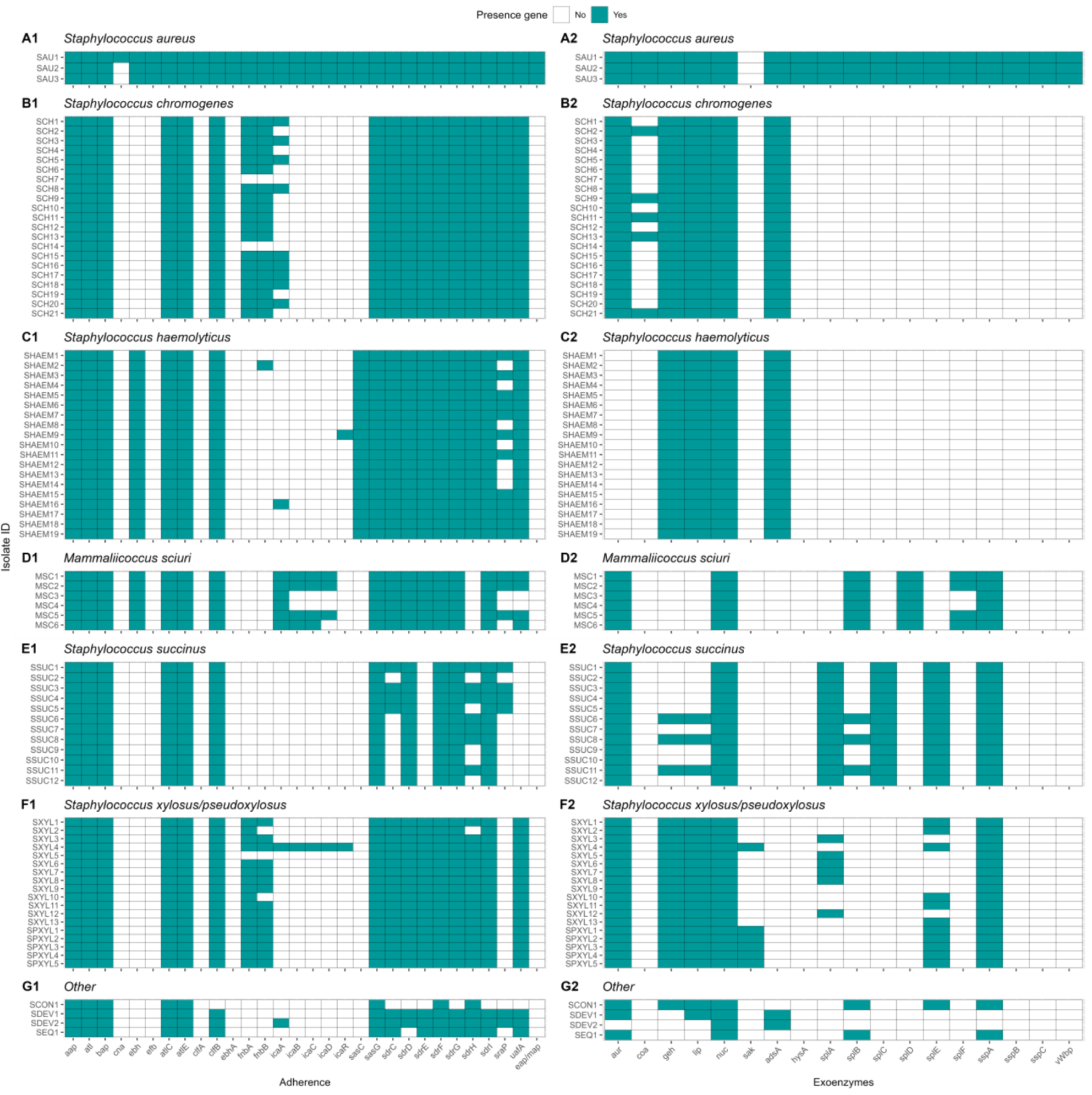
Distribution of virulence genes involved in adherence and exoenzymes in *Staphylococcus aureus* and non-*aureus staphylococci* and *mammaliicocci* (NASM). *Staphylococcus aureus* strain 1 (SAU 1) represents a reference strain: American Type Culture Collection strain Seattle 1945; *Staphylococcus aureus* subsp. *aureus* Rosenbach 25923 ^™^. Rosenbach 25923™. aap: accumulation associated protein. bap: biofilm associated surface protein. atl: autolysin. atlC: fibronectin binding autolysin. clfA-B: clumping factor class A-B. cna: collagen adhesin precursor. ebh: cell wall associated fibronectin-binding protein. ebhA: cell wall associated fibronectin-binding protein class A. efb: fibrinogen-binding protein. fnbA-B: fibronectin-binding protein class A-B. eap/map: extracellular adherence protein/MHC analogous protein. sasC-G: cell wall surface anchor family protein type C-G. sraP: serine rich adhesion for platelets. icaA-R: intercellular adhesion proteins class A-R. sdrC-I: Ser-Asp rich fibrinogen-binding bone sialoprotein-binding protein class C-I. uafA: cell wall anchored protein class A. adsA: adenosine synthase A. aur: aureolysin. coa: staphylocoagulase. geh: glycerol ester hydrolase. hysA: hyaluronate lyase. lip: triacylglycerol lipase. sak: staphylokinase. nuc: thermonuclease. spLA-F: serine proteinase class A-F. sspA-C: serine protease class A-C. vWbp: von Willebrand factor-binding protein.

**Figure 9.**
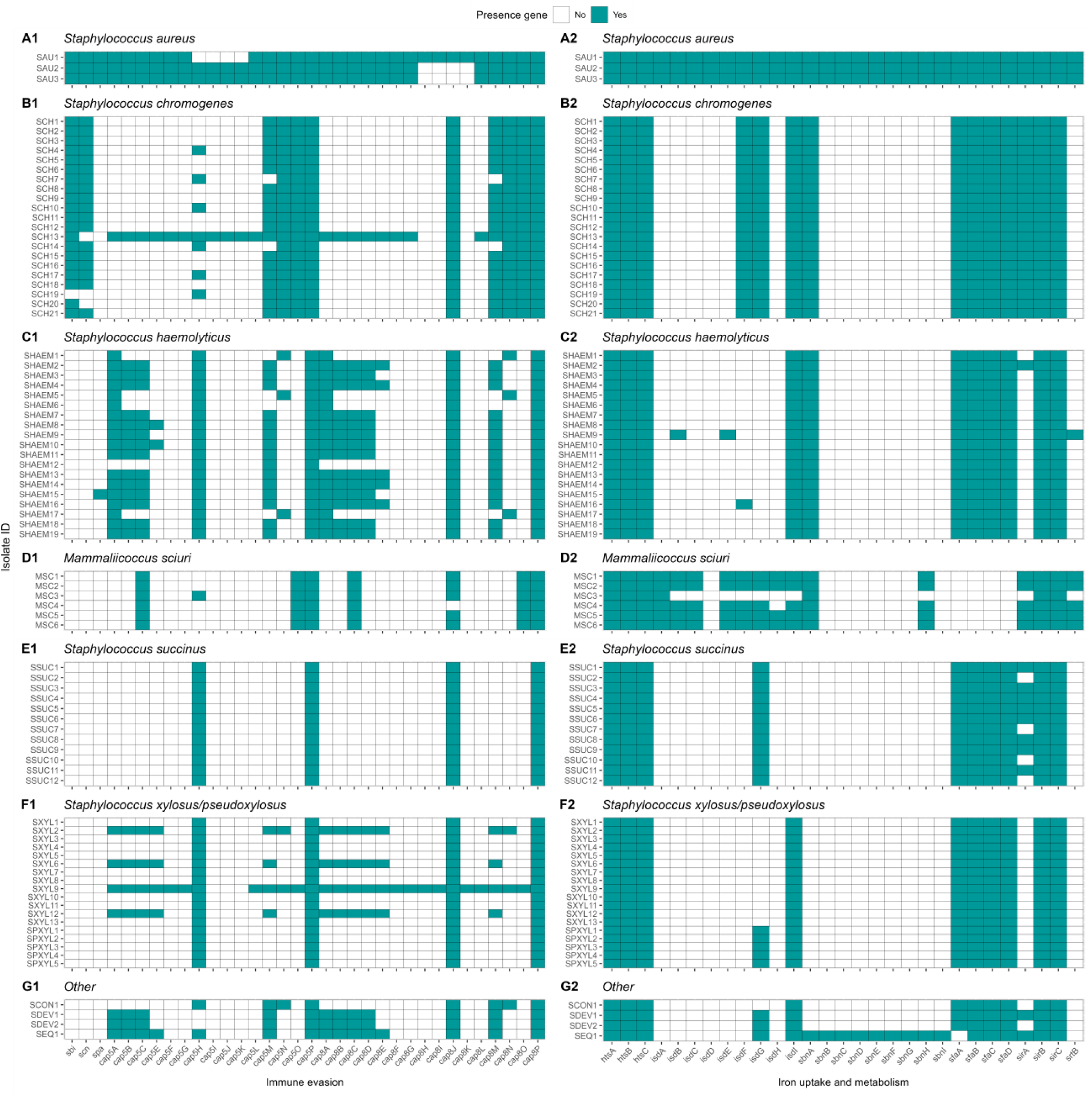
Distribution of virulence genes involved in immune evasion and iron uptake and metabolism in *Staphylococcus aureus* and non-*aureus staphylococci* and *mammaliicoci* (NASM). *Staphylococcus aureus* strain 1 (SAU 1) represents a reference strain: American Type Culture Collection strain Seattle 1945; *Staphylococcus aureus* subsp. *aureus* Rosenbach 25923 ^™^. cap8A-O: capsular polysaccharide-synthesis enzyme class 8A-O. cap5A-P: capsular polysaccharide-biosynthesis protein class 5A-P. chp: chemotaxis-inhibiting protein. sbi: staphylococcal binding immunoglobulin protein. scn: staphylococcal complement inhibitor. spa: staphylococcal protein A. isdA-I: iron-regulated surface determinant protein class A-I. htsA-C: FecCD iron compound ABC transporter permease family protein class A-C. sbnA-I: siderophore biosynthesis class A-I. srtB: sortase B. sirA-C: staphylococcal iron regulated class A-C. sfaA-D: staphyloferrin A biosynthesis protein class A-D.

**Figure 10.**
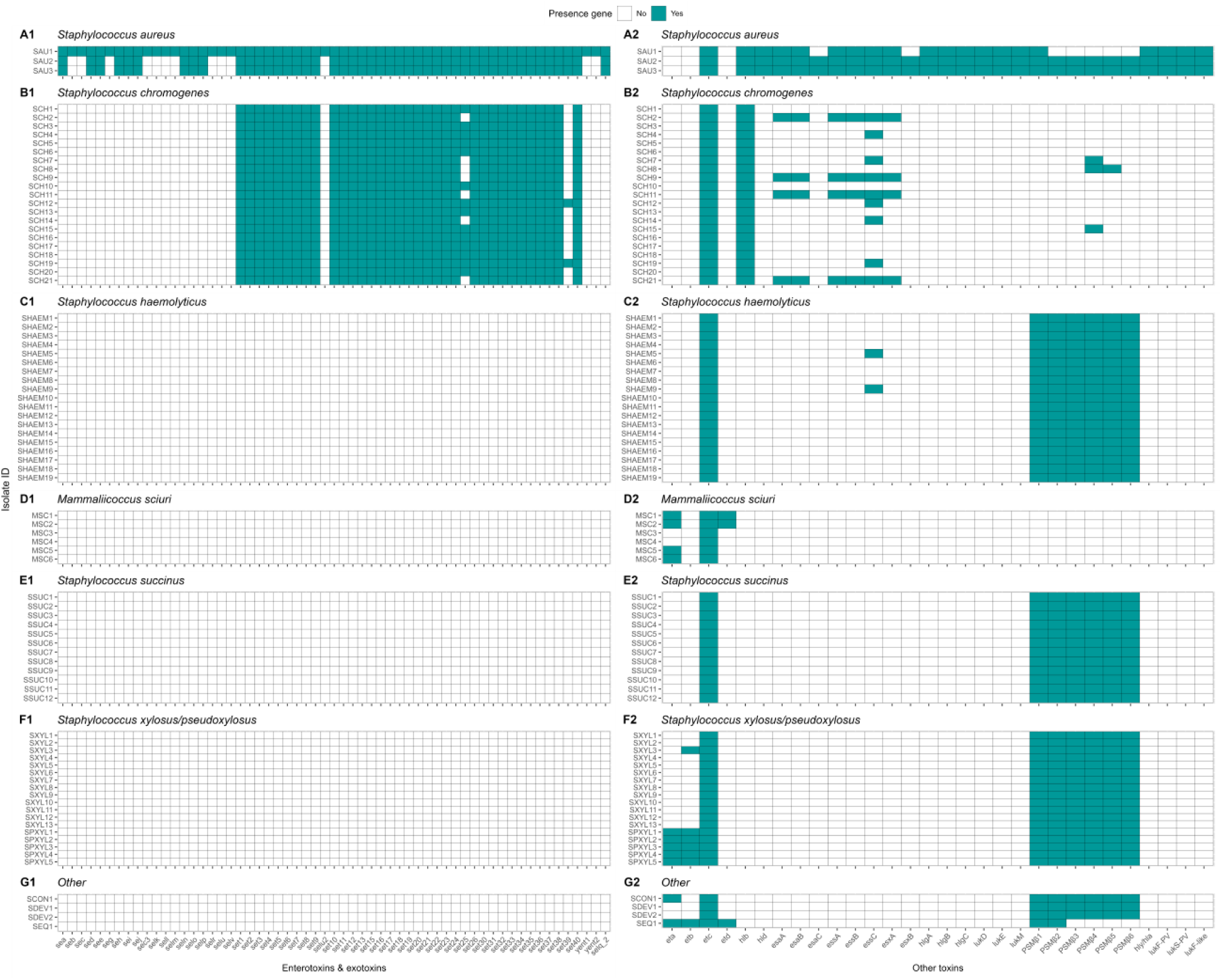
Distribution of toxin related genes in *Staphylococcus aureus* and non-*aureus staphylococci* and *mammaliicocci* (NASM). *Staphylococcus aureus* strain 1 (SAU 1) represents a reference strain: American Type Culture Collection strain Seattle 1945; *Staphylococcus aureus* subsp. *aureus* Rosenbach 25923 ^™^. se/sel a-v: Staphylococcal enterotoxin A-V. selu2: Staphylococcal enterotoxin U2. sec3: Staphylococcal enterotoxin C1 precursor. set1-40: staphylococcal exotoxin 1-40. yent1-2: enterotoxin yent1-2. hly/hla: alpha-hemolysin. hlb: beta-hemolysin. hld: delta-hemolysin. hlgA-C: gamma-hemolysin component A-C. lukD-M: leukocidin D-M. lukF-like: Panton-Valentine leukocidin LukF-PV chain precursor. lukF-PV: Panton-Valentine leukocidin chain F precursor. lukS-PV: Panton-Valentine leukocidin chain S precursor. tsst: toxic shock syndrome toxin-1. eta-d: exfoliative toxin class A-D. esaA-C: type VII secretion system protein class A-C. essA-C: type VII secretion system protein class A-C. esxA-B: type VII secretion system secreted protein class A-B. PSMβ1-6: phenol-soluble modulin beta class 1-6.

#### Genetic determinants of antimicrobial resistance

In this research, we identified the presence of 28 different ARGs (**Figure 11**).

**Figure 11.**
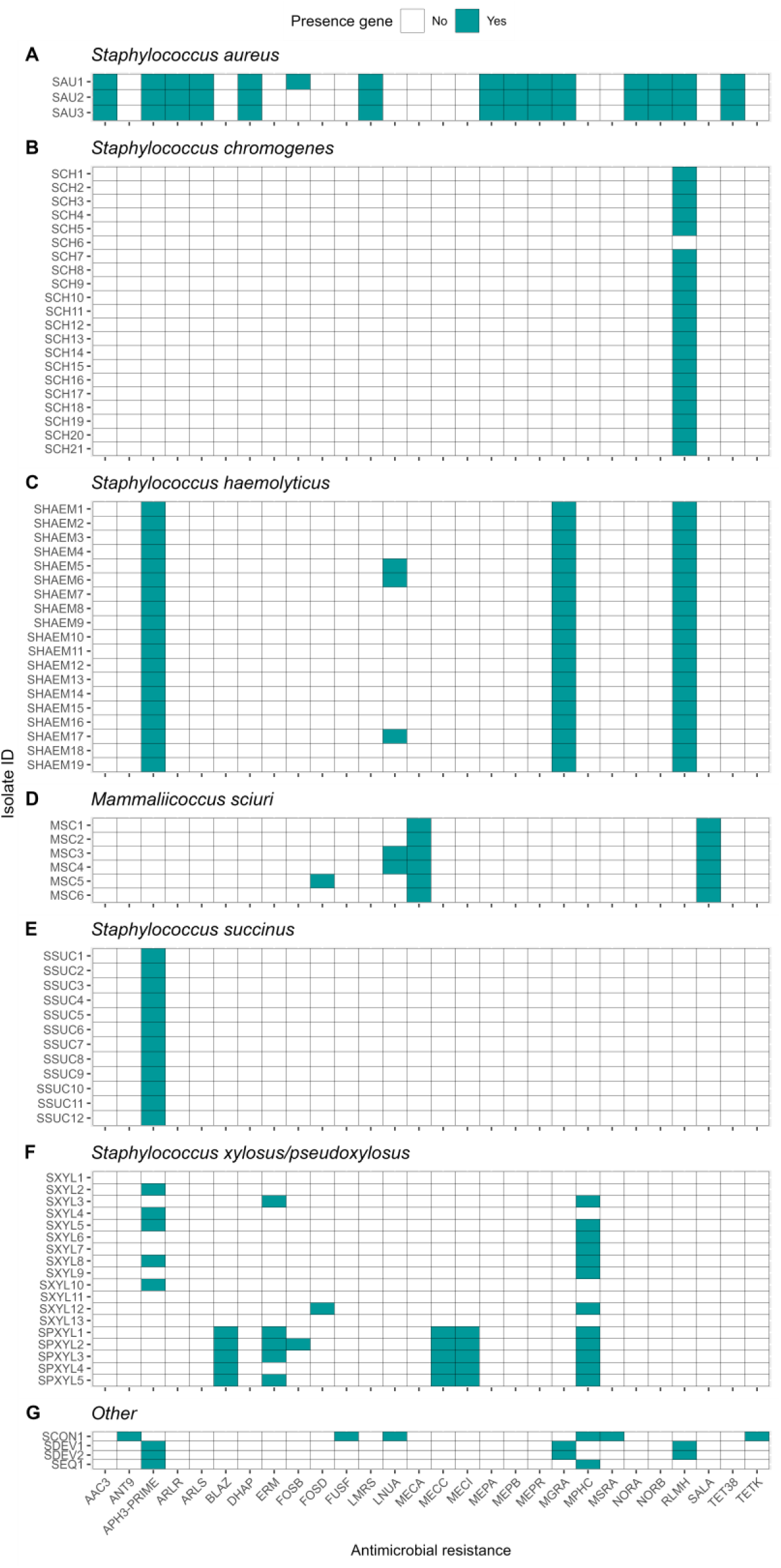
Distribution of antimicrobial resistance genes in *Staphylococcus aureus* and non-*aureus staphylococci* and *mammaliicoci* (NASM). *Staphylococcus aureus* strain 1 (SAU 1) represents a reference strain: American Type Culture Collection strain Seattle 1945; *Staphylococcus aureus* subsp. *aureus* Rosenbach 25923 ^™^. AAC3: Aminoglycoside N-acetyltransferase. ANT9: Aminoglycoside nucleotidyltransferase. APH3-PRIME: Aminoglycoside O-phosphotransferases. ArlR: Arl Response Regulator. ArlS: Arl sensor kinase. BLAZ: Class A betalactamases. DHAP: Phenicol resistance MFS efflux pumps. ERM: 23S rRNA methyltransferases. FOSB: Fosfomycin thiol transferases class B. FOSD: Fosfomycin thiol transferases class D. FUSF: Fusidic acid esterases. LMRS: Drug and biocide MFS efflux pumps. LNUA: Lincosamide nucleotidyltransferases. MECA: Penicillin binding protein class A. MECC: Penicillin binding protein class C. MECI: Penicillin binding protein class I. MEPA: Drug and biocide MATE efflux pumps class A. MEPB: Drug and biocide MATE efflux pumps class B. MEPR: Drug and biocide MATE efflux pumps class R. MGRA: MDR regulator. MPHC: Macrolide phosphotransferases. MSRA: MLS resistance ABC efflux pumps. NORA: Drug and biocide MFS efflux pumps class A. NORB: Drug and biocide MFS efflux pumps class B. RLMH: 23S rRNA methyltransferases. SALA: Multi-drug ABC efflux pumps. TET38: Tetracycline resistance MFS efflux pumps. TETK: Tetracycline resistance MFS efflux pump.

*Staphylococcus aureus* had 15 of these genes in its genome, whereas each NASM species had between 1 and 6 ARGs in at least one isolate. Ninety-seven percent of the isolates (81/83) showed the presence of at least 1 ARG. The most prevalent ARGs were 23S rRNA methyltransferases (**RLMH**) (53.0% [44/83]), involved in the synthesis of an rRNA methyltransferase related to resistance against lincosamides and macrolides. Another gene that exhibited high prevalence was aminoglycoside O-phosphotransferases (**APH3-PRIME**) (50.6% [42/83]), which encodes an aminoglycoside-modifying enzyme. Additionally, macrolide phosphotransferases (**MPHC**), a gene associated with macrolide resistance, was present in 16.9% (14/83) of the isolates. Other genes, including those linked to penicillin resistance (e.g., class A betalactamases [**blaZ gene**]) or methicillin resistance (e.g., penicillin binding proteins [**mecA**, **mecB**, **mecI**]), demonstrated low prevalence (<10%) among SAU and NASM isolates.

## DISCUSSION

This study improved our understanding of NASM genomes isolated from teat apices of organic dairy cows around parturition, with a focus on their species distribution, phylogeny, antimicrobial activity, virulence, and antimicrobial resistance. Multiple NASM species, and occasionally multiple strains within the same species, were identified on teat apices. NASM isolates ranked among the “top 10” with lowest MIC against SAU were more prevalent in cows without IMI compared to those with IMI. Although all NASM genomes contained at least one AMP, no association was found between presence of AMPs in NASM genomes and *in vitro* inhibitory activity against SAU or SUB. One clade within the SSUC group comprised 7 NASM isolates, with the majority demonstrating high *in vitro* antimicrobial activity. However, for other species, no association was found between clade membership and *in vitro* antimicrobial activity. Lastly, our results showed that although SAU genomes showed a higher prevalence of virulence genes and ARGs, all NASM contained at least some of these genes, with a distribution that appeared to be in most cases species-dependent.

### Relationship between teat apex colonization by *NASM,* in vitro inhibitory activity and presence of intramammary infections

The prevalence of NASM on teat apex was similar to that reported by other studies in conventional dairy farms (De Visscher et al., 2016; Mahmmod et al., 2018). Consistent with previous studies, we found a high prevalence of Staphylococcus chromogenes and observed differences in NASM species distribution across farms (De Visscher et al., 2016; Mahmmod et al., 2018). These differences have been previously attributed to differences in management practices across farms (De Buck et al., 2021) and are not surprising, given the differences in the prevalence and distribution of different microorganisms that cause IMI on organic dairy farms (Peña-Mosca et al., 2023).

The counts or the presence of NASM on teat apex were not associated with the presence of SAU or SSLO IMI. These results are inconsistent to those reported by prior studies, in which prepartum colonization of teat apices by NASM was related to a lower risk of postpartum IMI (Piepers et al., 2011) and subclinical mastitis (De Vliegher et al., 2003). These findings suggest that, in our study population, the mere presence of NASM on the teat apex did not appear to be a major factor influencing IMI risk. Other characteristics, such as the antimicrobial activity of NASM isolates, may be more relevant, as proposed by previous research (Reyher et al., 2012), and supported by studies reporting the in vitro antimicrobial activity of NASM isolates (Braem et al., 2014; Carson et al., 2017; Toledo-Silva et al., 2022).

We observed wide variation in the in vitro antimicrobial activity among NASM isolates. A considerable number of isolates showed no inhibitory activity at the maximum concentration tested, suggesting the absence of effective antimicrobial activity in these isolates. Moreover, we did not find evidence of a difference in the average inhibitory activity of NASM isolates against SAU or SUB between cows with and without an IMI. One possible explanation is that average antimicrobial activity may not be a major determinant of IMI risk, and that strong antimicrobial activity against specific pathogens may play a more important role in preventing infection (Nakatsuji et al., 2017). Supporting this idea, we found that transitioning dairy cows without an IMI were more likely to carry NASM isolates with the lowest MIC values against SAU compared to cows with an IMI. Although these findings should be interpreted with caution given the small sample size and wide confidence intervals, they suggest the potential for a protective effect of NASM isolates exhibiting strong *in vitro* antimicrobial activity against SAU on IMI risk by SAU or SSLO. Our findings are in line with human studies, where isolates with high phenotypic antimicrobial activity were commonly found on the skin of healthy individuals but were rarely identified in individuals with SAU skin infections (Nakatsuji et al., 2017). As a whole, our results suggest that commensal NASM may contribute to defense against mammary gland pathogen invasion, although further research is needed to confirm these observations.

### Relationship between *NASM taxonomy* and presence of in vitro antimicrobial activity

Nine NASM species were identified from teat apices of organic dairy cows. Consistent with previous studies, we observed important differences in antimicrobial activity across NASM species (Braem et al., 2014; Carson et al., 2017). Earlier studies using the cross-streak method reported antimicrobial activity in 9.1% to 13% of NASM isolates (Braem et al., 2014; Carson et al., 2017). In contrast, we used agar dilution methods (Wiegand et al., 2008), which aligns with the cross-streak method but provides quantitative measurements (Toledo-Silva et al., 2022).

To our knowledge, quantitative MIC data for NASM against mastitis pathogens, have only been reported for SCH, *Staphylococcus epidermidis*, and *Staphylococcus simulans*, with MIC values against SAU ranging from 5.10 to 8.24 log10 CFU/mL (Toledo-Silva et al., 2022). In our study, we observed variation in NASM inhibitory activity against SAU and SUB. *Staphylococcus haemolyticus* exhibited little to no inhibition, consistent with prior findings (Carson et al., 2017). In contrast, SSUC showed the strongest inhibitory activity, with nearly 50% of its isolates ranked among the "top 10" lowest MIC values. This proportion was notably higher than that of other species, where it did not exceed 20%, and differs from prior reports (Carson et al., 2017). These discrepancies may be explained by the observation that antimicrobial activity in previous human research appeared to be strain-specific rather than species-specific (Nakatsuji et al., 2017). Hence, it is plausible that the antimicrobial activity of NASM isolates is influenced more by strain-level variation rather than by species species.

### Association between the phylogeny, genotypic, and in vitro antimicrobial activity of NASM isolated from the teat apex

To investigate strain-level variation, we constructed five phylogenetic trees to examine the evolutionary relationships among the most common NASM species in this study. For most species, clade membership was not associated with IMI status or *in vitro* antimicrobial activity. In SSUC, however, we observed that isolates with the strongest *in vitro* inhibitory activity were concentrated within a single clade. In particular, one clade containing 7 genomes showed a lower MIC against both SAU and SUB compared to isolates in other clades. Additionally, all 5 isolates classified as part of the “top 10” group for highest inhibitory activity within SSUC were found in this clade, suggesting that specific genome features of these SSUC isolates may confer enhanced protection against SAU and SUB.

Our phylogenetic analysis revealed that a single cow could harbor multiple strains of a NASM species within a single sample (**Figures 2-5**; e.g., cow 171: SHAEM1 and SXYL1; cow 87835: SCH5 and SCH11). This finding aligns with previous research showing that the teat apex can be colonized by multiple strains of the same NASM species (Woudstra et al., 2023). Multi-strain infections have been documented in various human and animal pathogens, including *Staphylococcus* spp. (Balmer and Tanner, 2011) and similar patterns have been observed among mastitis-causing pathogens colonizing the mammary gland. Specifically multiple strains of SAU (Smith et al., 2005) or SUB (Zadoks et al., 2003) have been isolated from the mammary gland. However, evidence suggests that when multiple strains colonize the mammary gland, a single strain often dominates, as demonstrated with SUB (Pryor et al., 2009). The implications of multi-strain infections are still not well understood, but it has been hypothesized it could have an impact on disease dynamics (Balmer and Tanner, 2011). This could have important consequences for interpreting mastitis pathogen epidemiology and for decision-making on dairy farms (Exel et al., 2022). Previous studies have documented inter-strain differences in epidemiology (Zadoks et al., 2011), virulence potential (França et al., 2021; Monistero et al., 2020), and antimicrobial activity (Nakatsuji et al., 2017). These findings emphasize the importance of considering strain-level variation in study design. Methodological approaches that do not differentiate between strains and species may yield ambiguous or inaccurate results and could partly explain contradictory findings regarding NASM epidemiology (De Buck et al., 2021).

### Genotypic and in vitro antimicrobial activity

One intriguing observation was the ubiquitous presence of AMP-associated gene clusters in all examined genomes. This finding contradicts a prior study conducted with isolates from conventional dairies, reporting that 21.5% of isolates contained gene clusters associated with the production of AMPs (Carson et al., 2017). Several factors could explain this discrepancy, including updates to the databases used for gene annotation in recent years. Differences in methodology, such as the inclusion of additional steps to assess AMP gene cluster completeness (Carson et al., 2017), may also have contributed. Lastly, caution is warranted, as the presence of AMP-associated genes does not guarantee their expression or the production of functional peptides (Sánchez-Romero and Casadesús, 2020).

None of the detected AMP gene clusters were associated with *in vitro* antimicrobial activity. The prevalence of AIP class I was higher in cows without an IMI, whereas AIP class II was more prevalent in cows with an IMI, specifically among isolates classified as SCH. Auto-induced peptides are important for SAU pathogenesis, since they play an important role for the induction of expression of virulence factors through the accessory regulation system(Le and Otto, 2015; Wang and Muir, 2016). In NASM, the production of AIPs can repress the expression of this system in SAU, thus reducing the virulence potential of this pathogen (Canovas et al., 2016; Toledo-Silva et al., 2021). Extensive research has explored the use of AIP-mediated quorum sensing interference as a strategy for controlling SAU infections (Gray et al., 2013; Tan et al., 2018). Therefore, a higher prevalence of AIPs might be expected in cows without an IMI, consistent with the observed pattern for AIP class I in SCH, but not for AIP class II. Additional research is required to gain a deeper understanding of the potential applications of AIPs in the management of mastitis.

### Genetic determinants of virulence

We investigated 207 virulence genes involved in adherence, immune evasion, iron uptake, and toxin production. These genes were highly prevalent in SAU but less common among NASM. Adherence-related genes (i.e., aap and bap) were more prevalent than in previous studies (Fergestad et al., 2021; Naushad et al., 2019; Srednik et al., 2017; Tremblay et al., 2013). The IcaA gene, essential for biofilm formation, was detected in SCH and SSUC, aligning with earlier reports (Tremblay et al., 2013). Fibronectin binding factors (fnbA and fnbB), critical for SAU invasion (Campos et al., 2022), were nearly ubiquitous in SAU and also present in SCH, SXYL, and SPSXYL.

Short-chain dehydrogenase/reductase genes, key for *Staphylococcus* pathogenesis (França et al., 2021), were identified in all isolates. Genes encoding exoenzymes that facilitate host invasion were commonly detected (Tam and Torres, 2019). Nucleases and aur genes were found in all NASM isolates, similar to previous research (Fergestad et al., 2021; Naushad et al., 2019). Other genes exhibited variable prevalence. For example, adenosine synthase A was exclusive to SCH, SHAEM, and SDEV. The triacyglycerol lipase and glycerol ester hydrolase genes were highly prevalent in SCH, SHAEM, and SXYL but less common in SSUC and absent in MSC, consistent with previous findings (Naushad et al., 2019).

Capsular genes, which help bacteria resist phagocytosis (Cheung et al., 2021), were present in all isolates, as reported in previous studies (Fergestad et al., 2021; Naushad et al., 2019). Genes encoding staphylococcal binding immunoglobulin protein and staphylococcal complement inhibitor, both important for SAU immune evasion (Rooijakkers et al., 2005; Smith et al., 2011), were detected in SAU and SCH but not in other NASM species, aligning with prior research (Naushad et al., 2019).

Enterotoxins were identified only in SAU, contrasting with previous studies that reported them in NASM (Fergestad et al., 2021; Naushad et al., 2019). Staphylococcal exotoxins were only identified in SAU and SCH. This finding is interesting considering that SCH is the most prevalent NASM isolated from milk samples (De Buck et al., 2021). Some studies suggest SCH negatively impacts udder health (Fry et al., 2014; Supré et al., 2011; Valckenier et al., 2020), while others indicate a protective effect (De Vliegher et al., 2003). However, the presence of exotoxin genes in all SCH isolates raises questions about the extent of strain-level differences in virulence. Exfoliative toxin type C was prevalent in almost all NASM isolates, consistent with one prior study (Fergestad et al., 2021), but not with another investigation (Naushad et al., 2019). Beta phenol-soluble modulins (**PSMs**), important for pathogenesis, were widespread in most NASM except SCH and MSC. Intriguingly, alpha PSMs, which are considered highly potent, were absent in both SAU and NASM. Type VII secretion system proteins are important for virulence of SAU and other gram-positive pathogens (Spencer and Doran, 2022), and were primarily found in SCH, consistent with previous studies (Fergestad et al., 2021; Naushad et al., 2019).

### Genetic determinants of antimicrobial resistance

In this investigation, we identified that nearly all NASM isolates (97.7%) contained at least 1 ARG in their genome. In most cases, distribution of ARGs was species dependent; and if a given ARG was present within a species, it was typically present across all isolates in that species. The most prevalent ARG was *rlmH*, a gene that halts action of macrolides and lincosamides, whose identification was not reported in NASM in a prior study (Nobrega et al., 2018). In addition, the prevalence of *mphC*, another gene involved in resistance to macrolides, showed a 16.9% prevalence in our study, which is comparable to that of a prior study (Nobrega et al., 2018).

Half of the isolates carried the *aph(3’)-III* gene, encoding an aminoglycoside-inactivating enzyme, a much higher prevalence than reported in a previous study where these ARGs were nearly absent (Nobrega et al., 2018).

Penicillin resistance in SAU is typically mediated by beta-lactamase production (Nobrega et al., 2018). Previous studies have shown various degrees of penicillin resistance in both SAU and NASM isolates (Fergestad et al., 2021; Nobrega et al., 2018; Oliver and Murinda, 2012).

Methicillin resistance in SAU strains, mediated by acquisition of the staphylococcal chromosomal cassette containing *mec* genes, remains a major concern for both human and animal health(Holmes and Zadoks, 2011; McCarthy et al., 2012; Patel et al., 2021; Paterson et al., 2014). In our study, the prevalence of mec genes was low, with *mecA* detected only in SSC isolates. Previous studies also reported a low prevalence of *mecA* in MSC, although it has occasionally been identified in other NASM species (Fergestad et al., 2021; Nobrega et al., 2018).

### Internal validity

This study investigated the quantitative antimicrobial activity of NASM isolates against SAU and SUB. While this method represents a novel approach that has only recently been described for its use on mastitis pathogens, it allows for the investigation of more subtle variations in the inhibitory activity of bacteria (Toledo-Silva et al., 2022); however, it also has certain limitations.

For instance, the novelty of these methods in mastitis research, coupled with the absence of predetermined MIC cut-off points, complicates the interpretation of results. This makes it challenging to discern whether the absence or presence of inhibition is due to antimicrobial activity or merely a competition for nutrients that creates unfavorable conditions for the growth of target microorganisms in the media (De Vliegher et al., 2004b; Toledo-Silva et al., 2022). To overcome this limitation, we tested the self-inhibition of SAU and SUB, and we found that a considerable number of NASM isolates were able to inhibit the growth of these mastitis pathogens at lower concentrations than those necessary for self-inhibition of SAU and SUB, suggesting the presence of inhibitory activity. In addition, it is important to highlight that the utilized culture-based methods captured data from a few morphologically distinct isolates from each teat apex sample (De Visscher et al., 2016). These data were used to draw conclusions about the counts and prevalence of inhibitory NASM on the teat apex, which may not accurately represent the diversity or relative abundance of inhibitory NASM on the teat apex (Dean et al., 2024). This limitation is particularly relevant given that multiple species and strains can simultaneously colonize the teat apex, as was observed in our study.

In this investigation, steps to control for potential technical and biological confounders were taken. This included the randomization of samples for processing, matching by important confounders, and assessing the magnitude of confounding by including these variables in the models used in this study.

Nonetheless, it is vital to understand that some of our results could be still biased by other unmeasured confounders. For instance, cow-level factors that increased the likelihood of presence of IMI such as abnormally high or low body condition score, milk leakage (Fernandes et al., 2022), dysfunctional immune response (Sordillo, 2018), or dirty udders (Compton et al., 2007b; Schreiner and Ruegg, 2003) could have influenced our results. Moreover, the small sample size and resulting imprecision of estimates could hinder our ability to make inference from the results in this exploratory study. The use of composite milk samples represents a potential limitation of this observational study, considering the lower sensitivity of milk culture to detect IMI (Reyher and Dohoo, 2011; Toft et al., 2019). However, by culturing composite milk samples from each cow multiple times and defining IMI positivity as the presence of an IMI in at least one sample, we addressed this limitation, improving milk culture sensitivity for IMI detection (Buelow et al., 1996). Nevertheless, evaluating exposure and outcome at the cow-level poses a challenge, as it is possible that NASM with *in vitro* inhibitory activity could be present on the teat apex of one quarter while mastitis pathogens lead to an IMI in a different quarter, compromising the validity of this investigation (Bravata and Olkin, 2001).

Lastly, one major limitation of our study was the fact that almost two thirds of SAU and SSLO IMI were detected in the first postpartum milk sample. This suggests that IMI may have been acquired before calving. This finding is not surprising given the challenges to control mastitis without antibiotics (NMC, 2019) and the increased prevalence of SAU on organic dairy farms (Cicconi-Hogan et al., 2013; Pol and Ruegg, 2007). All the above limits our ability to determine if colonization of NASM with antimicrobial activity preceded the acquisition of SAU or SSLO IMI. Therefore, it is conceivable that other factors not explored in this study, such as the presence of a close-up pen with poor bedding management, high stocking density, high fly presence, and presence of IMI carried across lactations for multiparous cows, may be influencing the results and explaining why some of these animals calved with IMI (De Vliegher et al., 2012; Fernandes et al., 2022; Green et al., 2008).

### External validity

This study was conducted in two organic dairy farms located in Minnesota and Colorado. Considering the substantial differences in the prevalence NASM species across different dairy farms in both this and previous studies (De Visscher et al., 2016; Mahmmod et al., 2018), it is prudent to restrict the generalization of results to dairy farms managed under similar conditions.

In addition, because antimicrobial activity differed across different NASM species and even within the species, results should be confirmed including a larger number of isolates representative of different species that can be found on the mammary gland of dairy cows. Finally, to address the wide heterogeneity of NASM antimicrobial activity, strain-level identification might be warranted in future studies to further investigate bacterial clades and antimicrobial peptides associated with the presence of *in vitro* antimicrobial activity.

One important limitation of our approach is that we focused our investigation on DNA sequences, while it is known that the presence of a gene does not necessarily guarantee its expression or the production of a protein (Sánchez-Romero and Casadesús, 2020). Hence, it is plausible that this could explain some of the inconsistencies between the genotypes (e.g., presence of AMP gene clusters) and phenotypes (e.g., *in vitro* inhibitory activity). In addition, it is conceivable that the AMPs responsible for the observed *in vitro* antimicrobial activity against SAU or SUB in our study’s isolates have not been characterized, are not present in the utilized database (i.e., Bagel 4), or could not be identified due to bioinformatic challenges (e.g., the lack of homology in the genes encoding precursor peptides for AMPs) (Morton et al., 2015).

Another limitation of our study was the fact that in the majority of the dairy cows the IMI was already present on the first postpartum sample after calving. This limits our ability to assess the temporality between the presence of AMP gene clusters or *in vitro* inhibitory activity and the onset of new IMI by SAU or SSLO. Lastly, the low sample size, and small number of farms limits our ability to make inferences about the different NASM species, especially those with low prevalence (n<5). This is especially important, considering the substantial variation in the distribution of NASM species and strains across different farms. Therefore, results from this study should only be generalized to the NASM species identified in the current study and extrapolation to other species should be done with caution.

## CONCLUSIONS

Teat apices of organic dairy cows around parturition harbored multiple NASM species, and in some cases, multiple strains within the same species. NASM isolates classified among the "top 10" with the lowest MIC against SAU were more frequently found in cows without SAU or SSLO IMI compared to those with an IMI in early lactation. At least one AMP-associated gene was identified in all NASM isolates, indicating a high prevalence across species; however, their presence was not associated with *in vitro* (i.e., phenotypic) antimicrobial activity. The reconstructed phylogenetic tree for each NASM species contained multiple clades, indicating a non-clonal population that may reflect the population structure of the cows both within and across the two farms, and/or the evolutionary history of these bacteria within the two farms.

NASM in vitro antimicrobial activity differed across species but, apart from SSUC, was not associated with clade membership. Virulence genes were highly prevalent in SAU, but in most cases showed a low prevalence in NASM species, with a distribution that appeared to be species dependent.

## ACKNOWLEDGEMENTS

This study was funded by the Organic Agriculture Research and Extension Initiative (OREI) from the National Institute of Food and Agriculture (grant number: 2018-51300-28563). The first author, Felipe Peña-Mosca, was partially funded by Fulbright and Agencia Nacional de Investigación e Innovación from Uruguay (scholarship number: POS_FUL_2019_1_1008441). The authors thank the Laboratory for Udder Health at the University of Minnesota for performing milk cultures and for their assistance in taxonomic identification of isolates. We convey our appreciation to the University of Minnesota Genomics Core (UMGC) for their assistance in DNA extraction, library preparation, and sequencing support, as well as to the Minnesota Supercomputing Institute (MSI) for supplying the necessary resources for data storage and computation. We also want to express our deep gratitude to the numerous students, faculty, and staff for their assistance with sample collection and laboratory work. Additionally, we extend our thanks to the owners and managers of the herds who made this study possible. Felipe Peña-Mosca sampled in Minnesota, conducted laboratory work, cleaned and wrangled the data, conducted bioinformatics and statistical analysis, and prepared the initial and final draft of the manuscript. Tara Gaire conducted laboratory work and edited the manuscript. Chris Dean sampled in Minnesota, collaborated on bioinformatics, and edited the manuscript. Peter Ferm collaborated on bioinformatics and edited the manuscript. Pablo Pinedo conceptualized the study, coordinated sampling in Colorado, and edited the manuscript. Diego Manriquez sampled in Colorado and edited the manuscript. Noelle Noyes conceptualized the study, acquired funding, collaborated on bioinformatics, coordinated sampling and laboratory work, and edited the manuscript. Luciano Caixeta conceptualized the study, acquired funding, coordinated sampling and laboratory work, and edited the manuscript. The authors have not stated any conflicts of interest.

## DATA AVAILABITY

The raw sequence data generated in this study has been submitted to the Sequence Read Archive on NCBI (Accession: PRJNA1070794).

## Notes

### Competing Interest Statement

The authors have declared no competing interest.

### Summary of Updates

We have updated the manuscript to include whole genome sequencing data analysis, including phylogenetic analysis and the identification of virulence genes, antimicrobial resistance genes, and antimicrobial peptide genes. These additions substantially strengthen the manuscript by providing a more detailed genomic characterization of NASM isolates. Accordingly, we have added one coauthor who contributed significantly to the genomic data analysis component of the study.

